# Comparative genomics reveals multipartite genomes undergoing loss in the fungal endosymbiotic genus *Mycetohabitans*

**DOI:** 10.1101/2025.06.12.659383

**Authors:** Bhuwan Abbot, Sara Field, Lauren Carneal, Richard Allen White, Andra Buchan, Caroline West, Laira Lee, Morgan Elizabeth Carter

**Affiliations:** Department of Biological Sciences, The University of North Carolina at Charlotte, Charlotte, NC, 28223, USA; Department of Bioinformatics and Genomics, The University of North Carolina at Charlotte, Charlotte, NC, 28223, USA; CIPHER Center, The University of North Carolina at Charlotte, Charlotte, NC, 28223, USA; North Carolina School of Science and Mathematics – Morganton, Morganton, NC, 28655, USA

**Keywords:** pangenome, endofungal, transposable elements, chromid, Rhizopus, pseudogene

## Abstract

Endosymbiotic bacteria extensively impact phenotypes of their eukaryotic hosts, while experiencing dramatic changes to their own genome as they become more host-restricted in lifestyle. Understanding the trajectory of such a genome has largely been done through study of animal-associated bacteria, especially insect endosymbionts. Yet, endofungal bacteria provide another natural experimental model for investigating how microbial genomes change when living inside of a host cell. *Mycetohabitans* spp. are culturable bacterial endosymbionts of the Mucoromycota fungus *Rhizopus microsporus*. To investigate the genome dynamics resulting from the endohyphal nature of this emerging model genus, we long-read sequenced and assembled new complete genomes to combine with previous assemblies, creating a global dataset of 28 complete *Mycetohabitans* genomes. All genomes were between 3.3 and 3.9 Mbp in size and were multipartite, structured into two conserved replicons with some strains having an additional plasmid. Based on evolutionary rate and gene content analysis of the different replicons, we termed the two major ones a chromosome and chromid. The differential presence of a third, mobilome-rich plasmid in some strains and the proliferation of transposable elements provide putative mechanisms for recombination or gene loss. The conservation of intact prophage and putative toxin-antitoxin systems, and extensive enrichment of secondary metabolite clusters in the *Mycetohabitans* genomes highlight the dynamic nature of this reducing genome. With fungal-bacterial symbioses becoming increasingly apparent phenomena, lessons learned from this symbiosis will inform our understanding of bacterial adaptation to novel hosts, and the process of microbe-microbe coevolution.

**SIGNIFICANCE:** Genomes change when organisms become closely associated, as bacterial partners undergo less opportunities for genetic exchange and become more reliant on their hosts. Our pangenome analysis of the endofungal genus *Mycetohabitans* shows a snapshot of evolution where the bacteria are moving from a facultative towards an obligate relationship with their host fungi. We find that *Mycetohabitans* genomes are marked by transposable elements, pseudogenes, a large biosynthetic repertoire, and novel bacteriophage. We provide a genomic resource to match publicly accessible strains for future work on endosymbiotic genetics and evolution.

## INTRODUCTION

With the increasing ease of bacterial genome sequencing and metagenomics, genetic information on both culturable and unculturable microbes is becoming readily available. Consequently, bacterial evolution is now better understood to be a function of the organism’s lifestyle and environment, spotlighting endosymbiotic bacteria as particularly interesting cases to gain insight into genomic changes during coevolution (Wernegreen 2015; Keeling and McCutcheon 2017). It is widely understood through studies of insect symbionts, like the well-studied *Wolbachia* genus, that bacteria with obligate host relationships typically have undergone genome reduction, widespread gene loss, and rapid genome evolution as a result of AT richness (Wernegreen 2015; Porter and Sullivan 2023; Vancaester and Blaxter 2023).

Obligate endosymbionts only represent the segment in this evolutionary journey with high host dependence, and their intractability *in vitro* poses difficulties for investigating processes leading to such a reduced genome. In contrast, genomes belonging to diatom endosymbionts in a transitionary state have expanded and accumulated transposable elements and pseudogenes, compared to those in a more stable endosymbiosis that have undergone the expected TE and core gene loss (Grujcic, et al. 2025). Similarly, insect endosymbionts with larger genomes like those in the *Sodalis* genus have high pseudogenization (Boyd, et al. 2024). Clearly, studying genetic features in intracellular bacteria that are recently host-restricted is needed to reach broad conclusions about genomic evolution during endosymbiosis.

Diverse eukaryotes undergo bacterial colonization, from unicellular protists to multicellular organisms like animals, yielding symbiotic outcomes ranging from pathogenic to mutualistic. For example, a wide range of fungi are hosts to endosymbiotic bacteria varying in the permanence and co-dependence within relationships (Robinson, et al. 2021; Pawlowska 2024). Arbuscular mycorrhizal fungi (phylum: Glomeromycotina) are extensively found to have obligate relationships with endohyphal bacteria with highly reduced genomes, indicating a fairly ancient relationship (Naumann, et al. 2010). Multiple ascomycetes and basidiomycetes relevant to plant and human health have facultative bacterial endosymbionts impacting fungal phenotypes and environmental function, such as pathogenesis and metabolism (Steffan, et al. 2020; Moses and Carter 2025). However, not much is known about the mechanistic basis of bacterial adaptation to fungal hosts and few genomic studies have been undertaken for this niche (Amses, et al. 2023). Hence, fungal-bacterial endosymbioses represent a subfield with a wealth of untapped knowledge for new perspectives on genomic dynamics during bacterial evolution towards an endosymbiotic lifestyle.

A valuable emerging model for fungal-bacterial endosymbiosis is the interaction between the Mucoromycota fungus *Rhizopus microsporus* and *Mycetohabitans* spp. (Betaproteobacteria). Initial studies identified *Mycetohabitans rhizoxinica* (Mrh) as enabling the fungus to cause rice seedling blight, as the bacterium produces the antimitotic toxin rhizoxin needed for pathogenicity (Partida-Martinez and Hertweck 2005, 2007). Rm commonly occurs as cosmopolitan mold and is an opportunistic human and plant pathogen with a global distribution. Many Rm isolates have *M. rhizoxinica, M. endofungorum* (Mef), or unclassified *Mycetohabitans* spp. (Msp) residing within their hyphae (Lackner, et al. 2009; Dolatabadi, et al. 2016; Cabrera-Rangel, et al. 2022; Carpenter, et al. 2024). While not all Rm isolates harbor *Mycetohabitans* spp., the ones that do are dependent on their endosymbiont for fungal sporulation and are capable of only vegetive growth when cured (Partida-Martinez, et al. 2007; Mondo, et al. 2017). Formerly classified within the genus *Burkholderia* and considered part of *Burkholderia* sensu lato, *Mycetohabitans* spp. have a reduced genome size of about 3 Mbp compared to the 5-8 Mbp observed in free-living *Burkholderia*, suggesting a specialized symbiosis (Estrada-De Los Santos, et al. 2018). However, both partners can be cultured independently and the bacteria horizontally transferred, creating a tractable study system.

Because of their endosymbiotic nature, limited sampling of *Mycetohabitans* isolates and a lack of complete genome assemblies have been obstacles for comparative studies into the structural organization of the endosymbiotic genome and the genetic diversity within this genus. Yet, the symbiosis between Rm and *Mycetohabitans* spp. presents an opportunity to investigate the mechanisms and processes enabling genome loss, and consequently the evolution of an obligate endosymbiotic lifestyle at a transitional period. In this study, we present new complete genome assemblies via long-read sequencing and assess the pangenome of a global collection of 28 complete *Mycetohabitans* genomes. We compare evolutionary dynamics and genome composition of their replicons and investigate the secondary metabolite potential and phage presence within those genomes. Our results show an open pangenome with differential evolutionary stability and conservation across the multipartite genome, revealing the dynamics at play in a facultative-to-obligate transitional endosymbiotic genome.

## RESULTS

### Long read, complete genome sequences for *Mycetohabitans* strains

We screened *Rhizopus* spp. isolates from the Agricultural Research Service Culture Collection (NRRL) for the presence of endohyphal *Mycetohabitans* strains via PCR. We identified novel strains Mrh B10 and Mrh B203, and confirmed endosymbionts in strains expected to be hosts based on bacterial sequence present in previous short-read fungal genome sequencing data (Carpenter, et al. 2024). Positive cultures were used for extraction of the bacteria, and then genomic DNA preparation for Oxford Nanopore Technologies long-read sequencing. In total, we sequenced and assembled genomes for 20 bacterial strains (**Table 1, Table S1, Figure S1**), including Msp B46 which we re-sequenced due to our previous PacBio assembly not resolving into complete replicons (Carpenter, et al. 2024) and the type strain Mrh B1 which was previously available as short-read Illumina assembly (Lackner, et al. 2011). For re-sequenced strains, our new ONT assemblies had average nucleotide identity (ANI) values of greater than 99.97% to their respective previous assemblies, indicating robust base calling (Jain, et al. 2018).

**Table 1.** *Mycetohabitans* spp. genomes sequenced and assembled in this study.

| Bacterial strain | Genome size | Coverage | BUSCO score | %GC | # of replicons | Host accession | Nucleotide accession |
| --- | --- | --- | --- | --- | --- | --- | --- |
| *M. rhizoxinica* B2 | 3853378 | 108x | 99.4% | 59.59% | 3 | ATCC 20577 | CP176780-CP176782 |
| *Mycetohabitans* sp. B8 | 3404497 | 100x | 99.4% | 60.08% | 2 | CBS 308.87 | CP173344-CP173345 |
| *M. rhizoxinica* B10 | 3429290 | 108x | 99.4% | 60.07% | 2 | NRRL 2710 | CP176778-CP176779 |
| *M. rhizoxinica* B203 | 3429291 | 108x | 99.4% | 60.07% | 2 | NRRL 6203 | CP176776-CP176777 |
| *M. endofungorum* B23 | 3642467 | 111x | 99.2% | 59.76% | 3 | NRRL 62023 | CP184774-CP184776 |
| *M. endofungorum* B29 | 3355370 | 107x | 99.1% | 60.22% | 2 | NRRL 13129 | CP173342-CP173343 |
| *M. rhizoxinica* B34 | 3650605 | 101x | 99.3% | 59.97% | 2 | NRRL 2934 | CP173340-CP173341 |
| *Mycetohabitans* sp. B46 | 3327273 | 102x | 99.4% | 60.03% | 2 | NRRL 5546 | CP173320-CP173321 |
| *M. rhizoxinica* B48 | 3371874 | 100x | 99.2% | 59.92% | 2 | NRRL 5548 | CP173338-CP173339 |
| *M. rhizoxinica* B50 | 3532824 | 100x | 99.3% | 59.93% | 2 | NRRL 5550 | CP173336-CP173337 |
| *M. endofungorum* B51 | 3754811 | 105x | 99.6% | 59.47% | 3 | NRRL 5551 | CP173333-CP173335 |
| *M. rhizoxinica* B514 | 3853403 | 111x | 99.4% | 59.59% | 3 | NRRL 1514 | CP176773-CP176775 |
| *M. rhizoxinica* B52 | 3623805 | 105x | 99.1% | 59.89% | 2 | NRRL 5552 | CP173331-CP173332 |
| *M. endofungorum* B53 | 3608068 | 87x | 99.3% | 59.78% | 3 | NRRL 5553 | CP173328-CP173330 |
| *M. rhizoxinica* B55 | 3528370 | 99x | 99.4% | 59.87% | 3 | NRRL 6255 | CP173325-CP173327 |
| *M. rhizoxinica* B58 | 3668316 | 97x | 98.8% | 59.72% | 3 | NRRL 5558 | CP184771-CP184773 |
| *M. rhizoxinica* B62 | 3429197 | 116x | 99.4% | 60.07% | 2 | CBS 339.62 | CP178341-CP178342 |
| *M. endofungorum* B75 | 3642023 | 102x | 98.9% | 59.71% | 3 | NRRL 66675 | CP173322-CP173324 |
| *M. rhizoxinica* B82 | 3454439 | 102x | 99.1% | 59.92% | 2 | NRRL 2582 | CP184791-CP184792 |
| *M. rhizoxinica* B1 | 3749950 | 105x | 99.3% | 59.71% | 3 | ATCC 62417 | CP184277-CP184279 |

The genome sizes range from 3.3 Mbp to 3.8 Mbp (**Figure S2**), and all genomes were considered complete with BUSCO scores upward of 98.8% and having resolved into circular replicons (**Table 1**). The genomes all had lower than 0.6% contamination as reported via CheckM (Parks, et al. 2015). The GC content of all *Mycetohabitans* strains lies at ∼60% (**Table 1, Table S2**). All *Mycetohabitans* genomes have two conserved replicons – a chromosome (∼2.7 Mbp) and a chromid (∼700-900 kbp). A plasmid of ∼100-200 kbp is only found in 12 strains. The copy number of these plasmids within their bacterial hosts is not known, though they are related as they fall within the same MOB-Suite unique cluster, AG595 (**Table S3**) (Robertson and Nash 2018). For analyses in this study, we considered a pangenomic dataset of 28 total long read *Mycetohabitans* genomes (**Table S1**) comprised of these new ONT assemblies and our previous PacBio assemblies (Carpenter, et al. 2024).

### *Mycetohabitans* strains have atypical natural boundaries for average nucleotide identity

In recent genomic studies of *Burkholderia* sensu lato, *Mycetohabitans* strains were underrepresented due to the lack of available genomes (Mullins and Mahenthiralingam 2021; Bach, et al. 2022; Bach, et al. 2023). We used the Genome Taxonomy Database to phylogenetically analyze a representative set of *Burkholderia* sensu lato genomes. We also included those from closely-related *Mycoavidus* and *Glomeribacter* genera given their niche as endofungal bacteria and used *Pandoraea* as an outgroup (**Figure 1A**). The genus *Candidatus* Vallotia made up of bacterial symbionts of adelgids was previously proposed to be consolidated with *Mycetohabitans* (Dial, et al. 2022; Szabó, et al. 2022); *Candidatus* Vallotia strains are now listed in the Genome Taxonomy Database as *Mycetohabitans* and were thus included in our phylogenetic analysis. Based on the data in **Figure 1**, we have chosen to label them as *Candidatus* Vallotia and differentiate them from *Mycetohabitans*, further commented on in the discussion.

**Figure 1.**
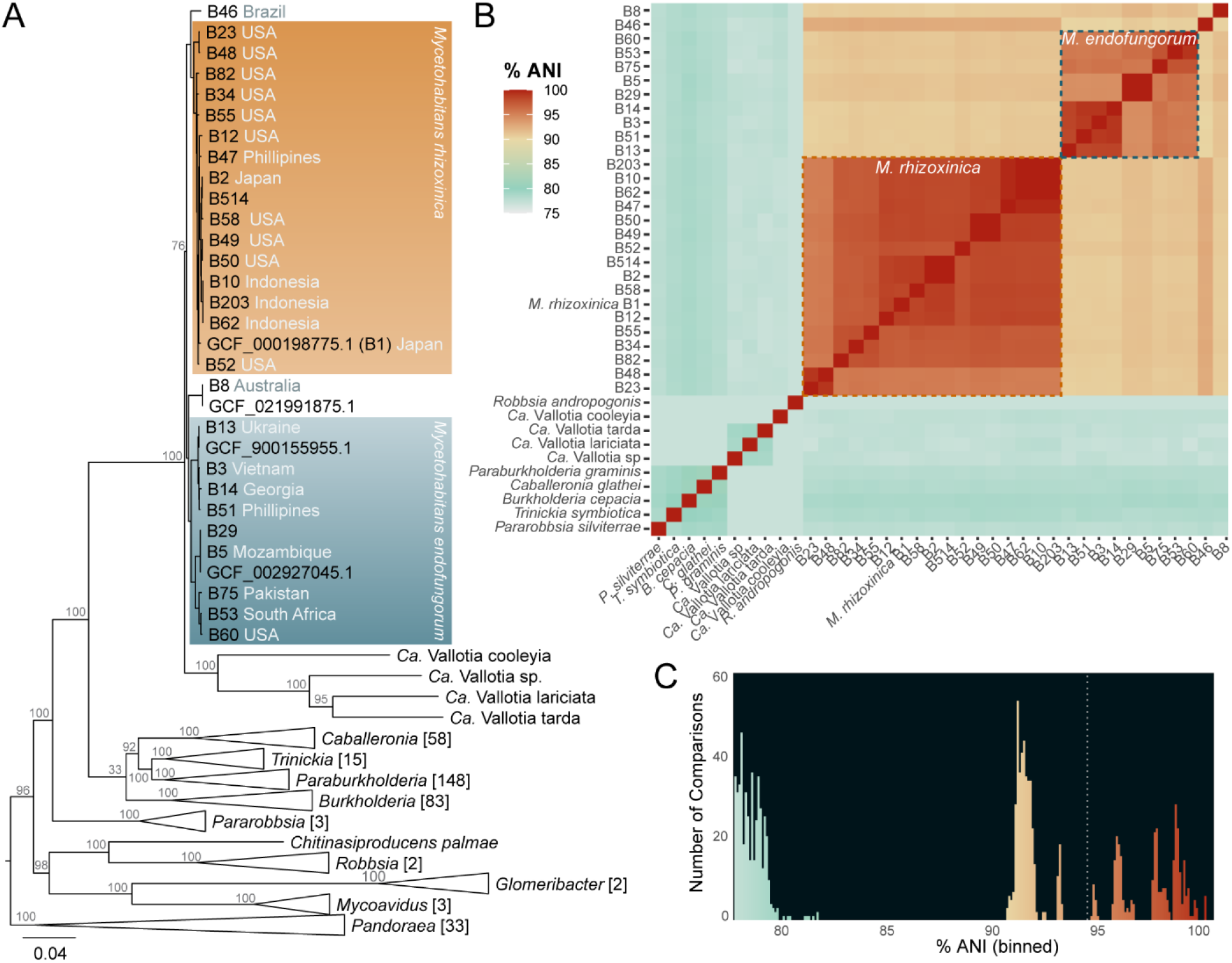
Taxonomic placement of *Mycetohabitans* strains within *Burkholderia* sensu lato. **A**. An approximately-maximum-likelihood tree for *Mycetohabitans* genomes and representatives from closely related genera provided by the Genome Taxonomy Database. Branches that contained a single genus were collapsed and the number of taxa within is displayed in brackets, except for the *Mycetohabitans* and *Ca*. Vallotia clades. Branch length indicates substitutions per site. When known, the country of origin is indicated for the host fungus of each *Mycetohabitans* strain (**Table S1**). Local support values were computed with the Shimodaira-Hasegawa test and are shown at relevant nodes (Shimodaira and Hasegawa 1999); all support values within *Mycetohabitans* spp. were >70 and are not shown. **B**. Average nucleotide identity (ANI) comparisons of the *Mycetohabitans* genomes to each other and representative strains from *Burkholderia* sensu lato (accession numbers in **Table S4**). Dotted lines enclose comparisons between the same species of *Mycetohabitans*. **C**. Count data for the ANI values in panel B binned by 0.1. A dotted line indicates the natural gap at 94%.

The *Mycetohabitans* genus forms a clade that is further validated by ANI analysis (**Figure 1B**). Inter-genus comparisons within *Burkholderia* sensu lato fall around 77-80%, a typical boundary for new genera. There is a discontinuity that begins around 83%, the expected beginning of the range for inter-species comparisons, but the intra-genus *Mycetohabitans* comparisons then span from 90% to >99% (**Figure 1C**). Comparisons between the type strain of Mef (B5) and Mrh strains are 90.7-91.9%, a rare range for ANI values (Jain, et al. 2018; Konstantinidis 2023). The orphan strains Msp B46 and Msp B8 share ∼92.5% ANI and ∼91.5% with Mrh and ∼90% and ∼90.5% with Mef, respectively. Mrh strains share >97% ANI, with the exception of B23 and B48 that are only 95.2% ANI with the other Mrh strains. Similarly, the type strain of Mef, B5, shares only 94.2-96.2% ANI with all but one strain (B29; >99% ANI). However, other strains within two Mef subclades share 95.5-96% ANI, such as B13 and B75 sharing 96% ANI, supporting that this is one species, with a potential subspecies structure (commented on further in the discussion).

### *Mycetohabitans* genomes show high synteny within replicons across strains

To investigate syntenic conservation across *Mycetohabitans*, we compared Mrh type strain B1 and Mef type strain B5 with the two isolates that did not fall into either named species, Msp B8 and Msp B46. Overall, protein cluster-based synteny between the four strains ranged from 69.9% to 80.8% (**Figure S3**). We found highly syntenic chromosomes, with somewhat less synteny observed in the chromids (**Figure 2A**). There were syntenic DNA blocks between the chromosome and chromid, most notably the chromid in Msp B8 has some elements syntenic with the Mrh B1 chromosome. In this analysis, Mrh B1 is the only strain that has a 3^rd^ replicon, the plasmid. The smaller syntenic blocks from the chromosomes or chromids of the other strains with the B1 plasmid include the loci for the ParB/RepB/Spo0J family partition protein (ACOALE_03960) and ParA family protein (ACOALE_03965) (**Table S5**). Additionally, we wanted to investigate the syntenic relationship between *Mycetohabitans* plasmids. Compared to that of the conserved replicons (chromosome and chromid), plasmids have lower shared protein sequence collinearity amongst each other, indicating higher diversity (**Figure S4**).

**Figure 2.**
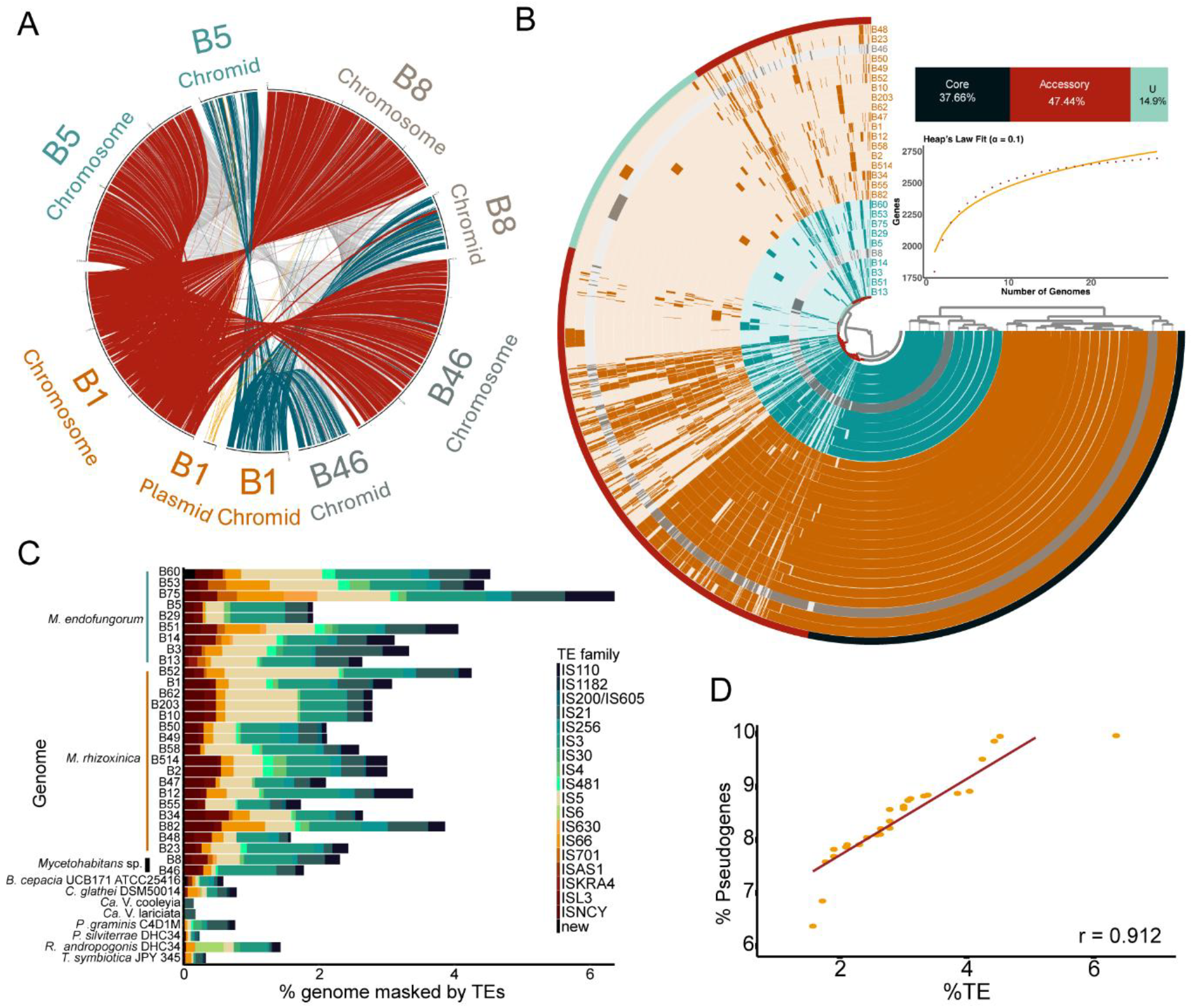
Genome synteny and content comparisons across the *Mycetohabitans* genus. **A**. Synteny comparison between replicons from the *Mycetohabitans rhizoxinica* (Mrh) B1, *Mycetohabitans endofungorum* (Mef) B5, *Mycetohabitans* sp. (Msp) B8, and Msp B46 genomes with B1 treated as the reference. Deep red connectors represent chromosome, deep blue chromids, and orange plasmid. **B**. Anvi’o plot illustrating the *Mycetohabitans* pangenome, with each ring representing one complete genome. The inner dendrogram shows hierarchical clustering of gene clusters. Core gene clusters (black), accessory gene clusters (deep red) and unique gene clusters (cyan) are marked as the outermost circle and their percentage in the pangenome is summarized in the stacked bar chart, below which is a Heap’s law fit. **C**. Stacked bar plot showing percent genome masked by transposable elements (TE). Columns are split by TE family. **D**. Correlation plot between percent of genome masked by TEs and percent of coding DNA sequences (CDS) that are pseudogenized. Pearson coefficient (r) is indicated at the bottom right of the plot. In all panels where relevant, *M. rhizoxinica* strains are indicated in orange, *M. endofungorum* in teal, and *M*. sp. in grey.

### The *Mycetohabitans* genus forms an open pangenome

To holistically explore the diversity of our 28 *Mycetohabitans* genomes, we generated an Anvi’o pangenome (**Figure 2B, Table S6**). 79,578 gene calls were clustered into 4,652 gene clusters which were further categorized as single-copy core genes (SCG), core, accessory and unique. While the SCGs were characterized by the program, core, accessory and singleton GCs were manually curated based on presence of the GC in all 28 genomes, 2 to 27 genomes, or only 1 genome, respectively. There were 1,752 core gene clusters with 52,055 gene calls (37.6%), 2,207 accessory gene clusters with 26,814 gene calls (47.44%) and 693 singleton gene clusters with 707 gene calls (14.89%).

The categories cell wall mobility, translation, transcription, mobilome, signal transduction and defense mechanisms have high % gene presence in the *Mycetohabitans* genome (**Figure S5**). There is no significant difference in COG percentages between the two species Mef and Mrh, except for the cell division category which was higher in Mef (**Figure S6**). A Heap’s Law model (Tettelin, et al. 2008) was used to estimate whether the *Mycetohabitans* pangenome is open or closed. Since the number of new genes increases upon addition of more genomes (as indicated by the α of 0.1), the *Mycetohabitans* genus forms an open pangenome (**Figure 2B**).

### *Mycetohabitans* strains have high pseudogene and transposable element content

To investigate the pseudogene and transposable element (TE) content in *Mycetohabitans* genomes, we used the PGAP annotations for each of the genome, and the publicly available annotations for representative *Burkholderia* sensu lato endosymbionts listed in **Table S4**. We computed the percentage of pseudogenized coding gene sequences (CDS) that are frameshifted, incomplete, or contain internal stop codons (Tatusova, et al. 2016). The total putative pseudogene content ranged from 6.4% in Mrh B47 genome to 9.96% in Mef B13. Strains with the longest plasmids - Mef B14, B75, and B51 – of lengths 201,412, 181,953, and 231,357 respectively have 9.95%, 9.86% and 9.56% of their CDS putatively pseudogenized (**Figure 2D, Figure S7, Table S5**). However, this pattern of correlation between plasmid length or presence, and pseudogene content is not consistent across *Bukholderia* sensu lato, as symbionts with other host ranges had pseudogene content ranging from 3.2% to 5.9% (**Figure S7, Table S5**).

Our analysis of TEs found that Mef B75 had the highest percentage (6.36%) of its genome masked by TEs, and Mrh B48 had the least, 1.58% (**Figure 2C, Table S5**). The highest fraction of TE families in *Mycetohabitans* were IS5 and IS30. Compared to other symbiotic *Burkholderia* sensu lato genomes, *Mycetohabitans* spp. have much greater portions of their genomes masked by TEs as well as a greater number of insertion elements (**Figure S8**). For example, *Robbsia andropogonis* had only 1.43% of its genome masked by TEs, lower than the least of *Mycetohabitans*, with IS4 as the family occupying the highest percent (**Figure 2D**). *Candidatus* Vallotia strains were found to have one IS21 element on their genome, masking only 0.2% of their genomes (**Figure 2D, Figure S8, Table S5**).

We wanted to check whether there is a correlation between TE and pseudogene content across the *Mycetohabitans* strains. Mef B75 and B51 both fall at the higher end of TE and pseudogene content, Msp B46, Mrh B47, B48, B49 and B50 all have both – low TE and pseudogene content (**Table S7, Figure 2C**). Interestingly Mef B60 has the second highest TE content of 4.53%, and falls towards the lower end of pseudogene content (**Table S7, Figure S7-8**).

### All *Mycetohabitans* genomes have a secondary core replicon

Though originally described as a megaplasmid in the Mrh B1 reference genome announcement, the presence of the secondary replicon and its conservation across *Mycetohabitans* strains supports a more stable role as a chromid (Harrison, et al. 2010; Lackner, et al. 2011). To assess the characteristics of the chromid and structural makeup of the pangenome, we created single replicon “pangenome” comparisons (Eren, et al. 2020). Anvi’o identified 3,500 and 1,335 gene clusters in the chromosomes and chromids, respectively (**Figure 3A-B**). The hierarchical clustering based on gene cluster frequencies within the two replicons group the strains similarly, with the exception that Msp B8 changes positions significantly by clustering with Mrh chromosomes, but Mef chromids. However, upon comparing the replication initiation protein-based phylogeny between the two, there are no significant signs of horizontal transfer of the chromids, indicating the observed change in clustering is likely due to a lack of genomes similar to B8 (**Figure S9**). The chromosome has a higher fraction of core gene clusters compared to the chromids which are conversely more enriched in accessory gene content (**Figure 3A-B**). Additionally, the chromids lack the chromosome replication initiator protein DnaA, and instead have a separate replication initiation protein and partitioning-related proteins similar to that of the *Mycetohabitans* plasmids, i.e. the ParA/B family proteins (**Table S6**). We used the tool Infernal to find the presence and location of rRNA or tRNA loci on the chromids but only found the presence of catalytic introns, riboswitches and C4 antisense RNAs. However, chromids do have some critical genes including the 30S ribosomal protein S21.

**Figure 3.**
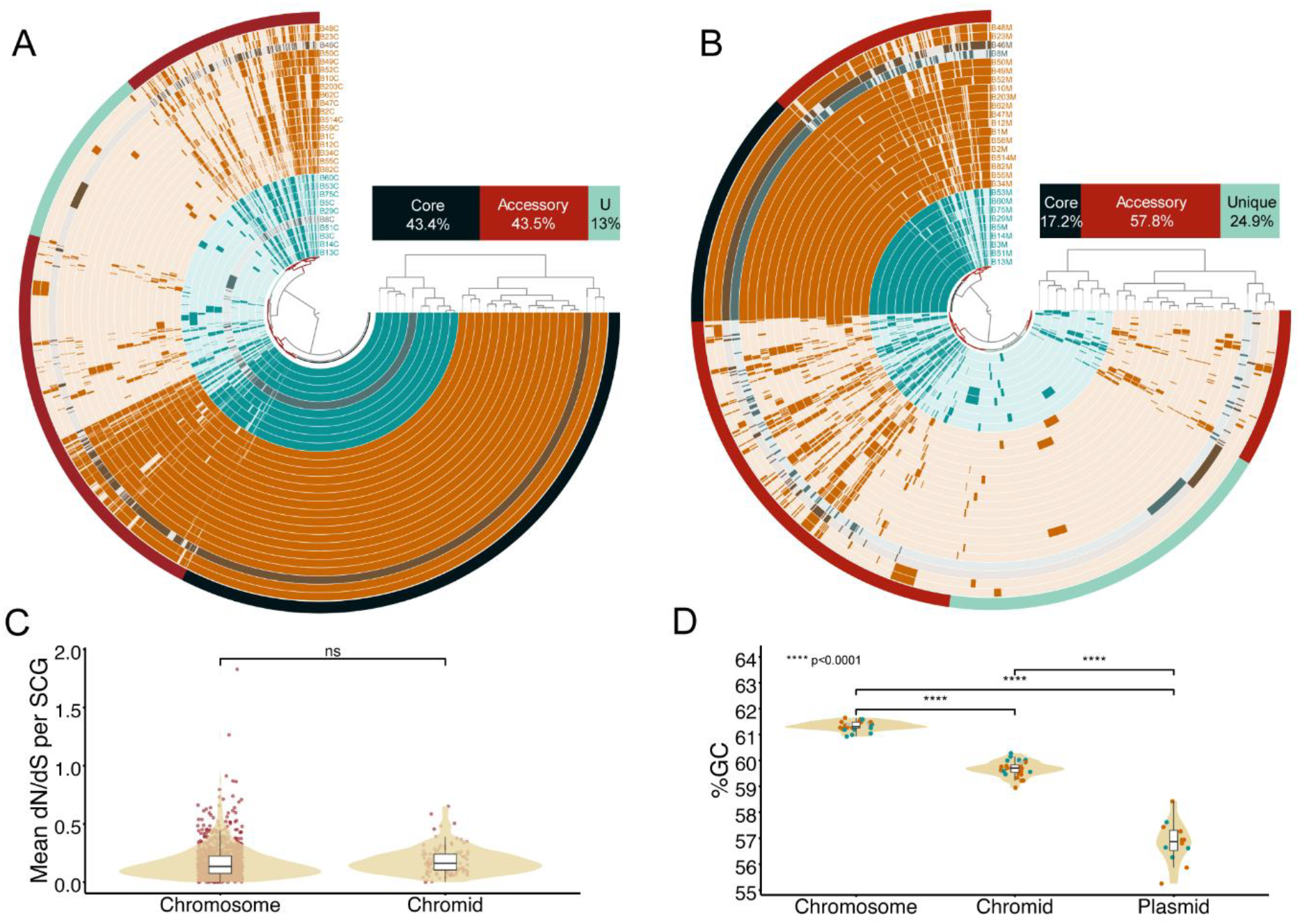
Comparisons between replicons i.e. chromosome, chromid and plasmid across *Mycetohabitans*. Anvi’o plots to compare. **A**.chromosomes and **B**. chromids across the strains. Each ring represents one *Mycetohabitans* strain’s relevant replicon and the inner dendrogram shows hierarchical clustering of gene clusters. Core gene clusters (black), accessory gene clusters (deep red) and unique gene clusters (cyan) are marked as the outermost circle and their percentage in the pangenome is summarized in the stacked bar chart. **C**. dN/dS value distribution for non-identical single-copy core genes in chromosomes and chromids **D**. Guanine-cytosine (%GC) content of each replicon is shown as a percentage with individual data points colored by species. P-value notations determined by T-test are indicated on the plot as asterisks. In all panels where relevant, *M. rhizoxinica* strains are indicated in orange, *M. endofungorum* in teal, and *M*. sp. in grey.

As the differential amount of gene cluster conservation suggests differential rates of evolution, we estimated the accumulation of non-synonymous (dN) to synonymous (dS) mutations in these replicons (Kryazhimskiy and Plotkin 2008). We calculated the dN/dS values for single-copy core genes in all *Mycetohabitans* strains that had non-identical sequences to B1, and plotted all those data points for each of the two replicons (**Figure 3C**). While greater dN/dS ratio indicates diversifying selection or adaptive evolution, a lower value would support higher selective or purifying forces acting on the replicon. The dN/dS ratio of chromid genes distributes similarly to that of the chromosome, supporting the description as a chromid and not a megaplasmid (Harrison, et al. 2010). We further explored this distinction by comparing the guanine-cytosine content (%GC) between chromids and chromosomes of the same strain. The %GC of the chromid is on average 1.65% lower than that of the chromosome (difference ranging from 0.9%-2.7%), and %GC of the plasmid is on average 4.43% lower than that of the chromosome, ranging from 2.83%-6.05% (**Figure 3D; Table S2**). Though the standard chromid definition requires the %GC of a chromid be within 1% of a chromosome, the *Mycetohabitans* chromid %GC are more conserved and much closer to that of the chromosomes versus the inconsistent third replicon (Harrison, et al. 2010).

In conclusion, we have termed the secondary replicon as a chromid due to (1) being the second largest replicon, (2) having chromosome-like nucleotide composition, (3) having plasmid-like replication systems, (4) having evolutionary rate similar to that of the chromosome, and (5) containing essential genes such as cell wall mobility, signal transduction and transcription (**Figure S5**).

### *Mycetohabitans* plasmids are dynamic accessory gene hubs that are potentially lost or transferred

We used MOB Suite to characterize the plasmids found in *Mycetohabitans* strains, and predict their mobility and host range (Robertson and Nash 2018). Overall, the predicted replicon type of the plasmid is not known. All but Mrh B23 plasmid have a relaxase belonging to MOBF family. Mrh B1 has an additional MOBP type relaxase, which may be why is has a putatively larger predicted host range of Pseudomonata. The predicted host range for the rest of the plasmids is within order Burkholderiales, with the exception of Mrh B23 which could not be predicted; the plasmids found in all strains except Mrh B23 are predicted to have conjugative mobility with a mating pair formation (MPF) type F. Mrh B23 having the shortest plasmid, and lacking a putative relaxases and mobilization mechanisms, suggests that it may be an artifactual remnant as a result of genome loss in this endosymbiont (**Table S3**).

A similar comparative analysis of the 12 *Mycetohabitans* plasmids using Anvi’o identified 362 gene clusters: 8 core, 218 accessory, and 136 unique (**Figure 4A**). The largest plasmid, Mef B51, has the highest number of genes and unique gene clusters. Out of the eight core gene clusters, only four are classified as single-copy core genes (SCGs) (Parks, et al. 2018), emphasizing the highly accessory nature of plasmid genes. Two of these SCGs are components of Type II toxin/antitoxin system – RelE/ParE family toxin, and RelB family antitoxin, with another being unannotated and the final being Replication Protein O. We binned plasmid gene clusters into mobilome-related components and found 45 gene clusters related to transposases, 55 related to toxin-antitoxin system, 20 related to conjugation machinery/Type IV Secretion System, and 2 related to bacteriophage genome fragments (**Figure 4A**). Conjugation-related proteins are a conserved feature of all *Mycetohabitans* plasmids except for Mrh B23; only one gene cluster related to conjugation was found in the plasmid of Mrh B23, the pilus assembly gene, *hicB*, that can also be found in B55.

**Figure 4.**
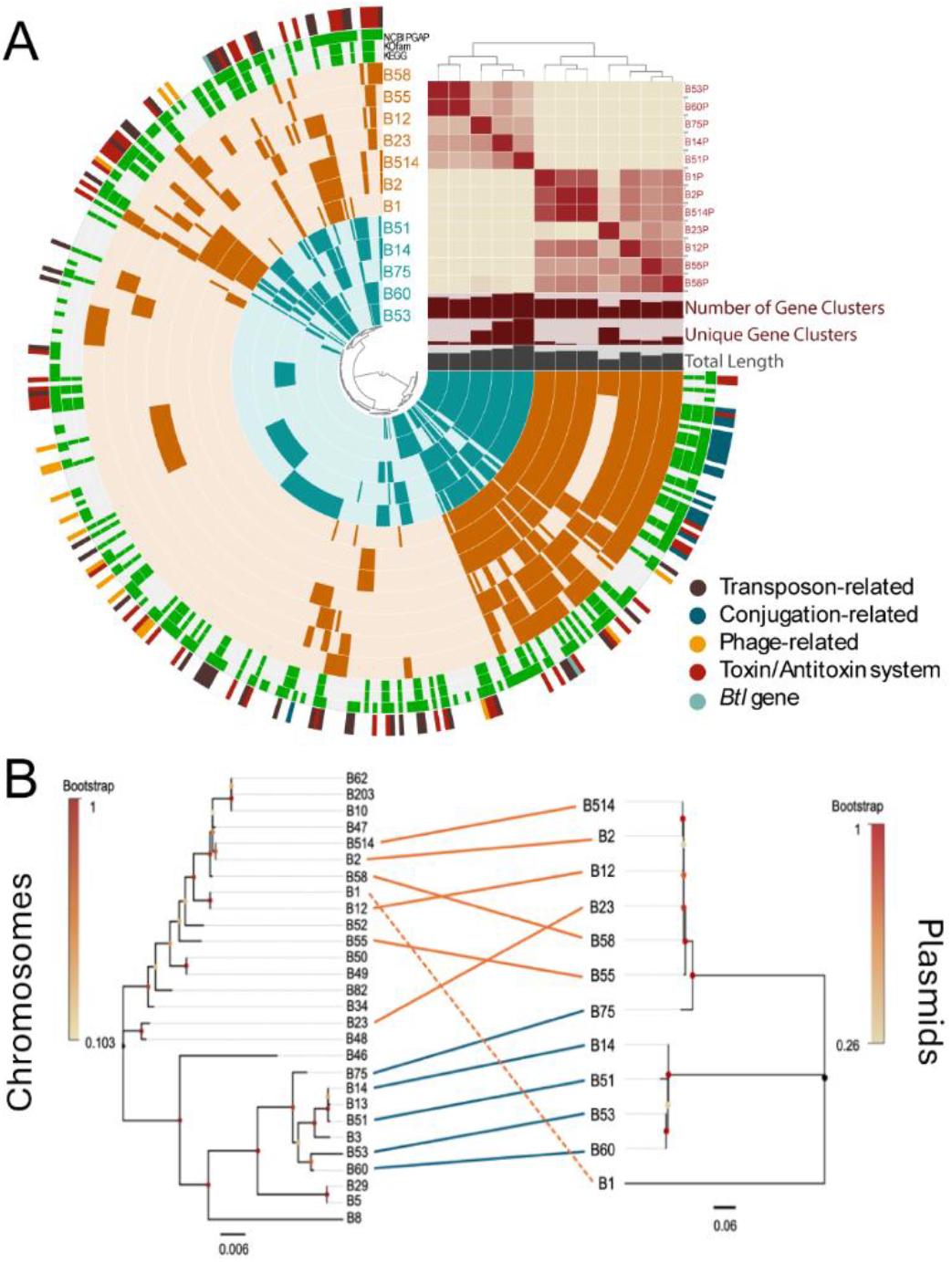
Analysis of the inconsistently present third replicon across *Mycetohabitans* strains. **A**. Anvi’o plot of *Mycetohabitans* plasmids indicated as individual rings, with toxin/antitoxin system, *btl* type III effector gene, and mobilome-related gene clusters annotated on the outermost circle colored according the panel key. Average nucleotide identity (ANI) of the plasmids is depicted as a heat map. Bar plots show the total lengths and number of gene clusters (unique and overall) for each plasmid sequence. The inner dendrogram shows hierarchical clustering of gene clusters. **B**. Co-phylogeny comparing topology of *dnaA* relationship across chromosomes (left) to *parA* relationship across plasmids (right) inferred by Maximum Likelihood with MEGAX. Dots at inner nodes of trees represent bootstrap values as a fraction of 1000 bootstraps. In all panels where relevant, *M. rhizoxinica* strains are indicated in orange, *M. endofungorum* in teal, and *M*. sp. in black.

One of the accessory gene clusters on the plasmid including strains Mrh B12, Mef B14, Mrh B1, Mrh B2, Mrh B514, Mef B51, Mef B53, Mrh B58, and Mef B75 contains *btl* genes which are known type III secretion system effector proteins that vary in number, sequence, and genomic location between strains (Carter, et al. 2020; Carpenter, et al. 2024). Overall, the *btl* genes are found across all three replicons and vary in number of copies for different strains, consistent with previous findings (Carpenter, et al. 2024). As no other effector proteins are validated, we did not further explore the diversity or genomic location of type III effectors.

To check the conservation of plasmids within bacterial lineages and assess the possibility of horizontal exchange, we created phylogenetic trees based on core replication genes: the *parA* gene found on plasmid and the *dnaA* gene on the chromosome (**Figure 4B**). Within Mef, plasmid presence and relatedness mirrors the chromosome topology and is monophyletic. Yet, plasmid loss likely occurred in strains such as Mef B13 and B3 which cluster with plasmid-containing strains despite lacking plasmids themselves. However, within Mrh, there is more discord, as which strains do or do not have a plasmid is not as clearly clustered. Mrh B1 stands out with the most divergent plasmid from other Mrh strains, contrasting with the strain’s overall close relatedness with Mrh B12 (**Figures 1-2**). That Mrh B12 and B1 have such distinct plasmids suggests plasmid horizontal transfer has occurred within the Mrh lineage, a hypothesis that is further supported by B23 and B58 having similar plasmids despite their overall divergence (**Figure 4B**).

### *Mycetohabitans* spp. are enriched in diverse biosynthetic gene clusters, with many associated with nonribosomal peptides

From previous studies, it is known that *Mycetohabitans* spp. have a high percentage of their genome devoted to biosynthetic gene clusters (BGCs) compared to other species in *Burkholderia* sensu lato (Mullins and Mahenthiralingam 2021). We identified a total of 373 unique biosynthetic gene clusters (BGCs) across the 28 *Mycetohabitans* strains using antiSMASH (**Figure S10**) (Blin, et al. 2023). On average, there are 13.3 BGCs per strain and no significant difference in the number of BGCs per species (anova, p<0.05). All strains were predicted to have BGC clusters for a terpene, a terpene/phosphonate, ribosomally synthesized lassopeptides, and numerous distinct nonribosomally synthesized peptides (NRPs) (**Figure 5A**). The majority of BGCs were found in chromosomes and chromids, with only two clusters predicted in plasmids: NRP group 29 in Mef B51 and NRP group 27 in Mef B75 (**Figure 5B; Table S8**). The classic rhizoxin cluster that led to the discovery of this genus was putatively found in all strains except Mrh B82 (Partida-Martinez and Hertweck 2005, 2007). B8 was the only strain predicted to have a hybrid polyketide synthase (PKS) and NRPS cluster other than rhizoxin, which produces necroxime (Niehs, et al. 2020). The conservation of predicted ribosomally synthesized lassopeptide BGCs align with previous findings on a smaller set of genomes showing loci with homology to those from type strain B1 that produce burhizin-23, mycetohabin-16, and mycetohabin-15, though their ecological role remains unknown (Bratovanov, et al. 2020).

**Figure 5.**
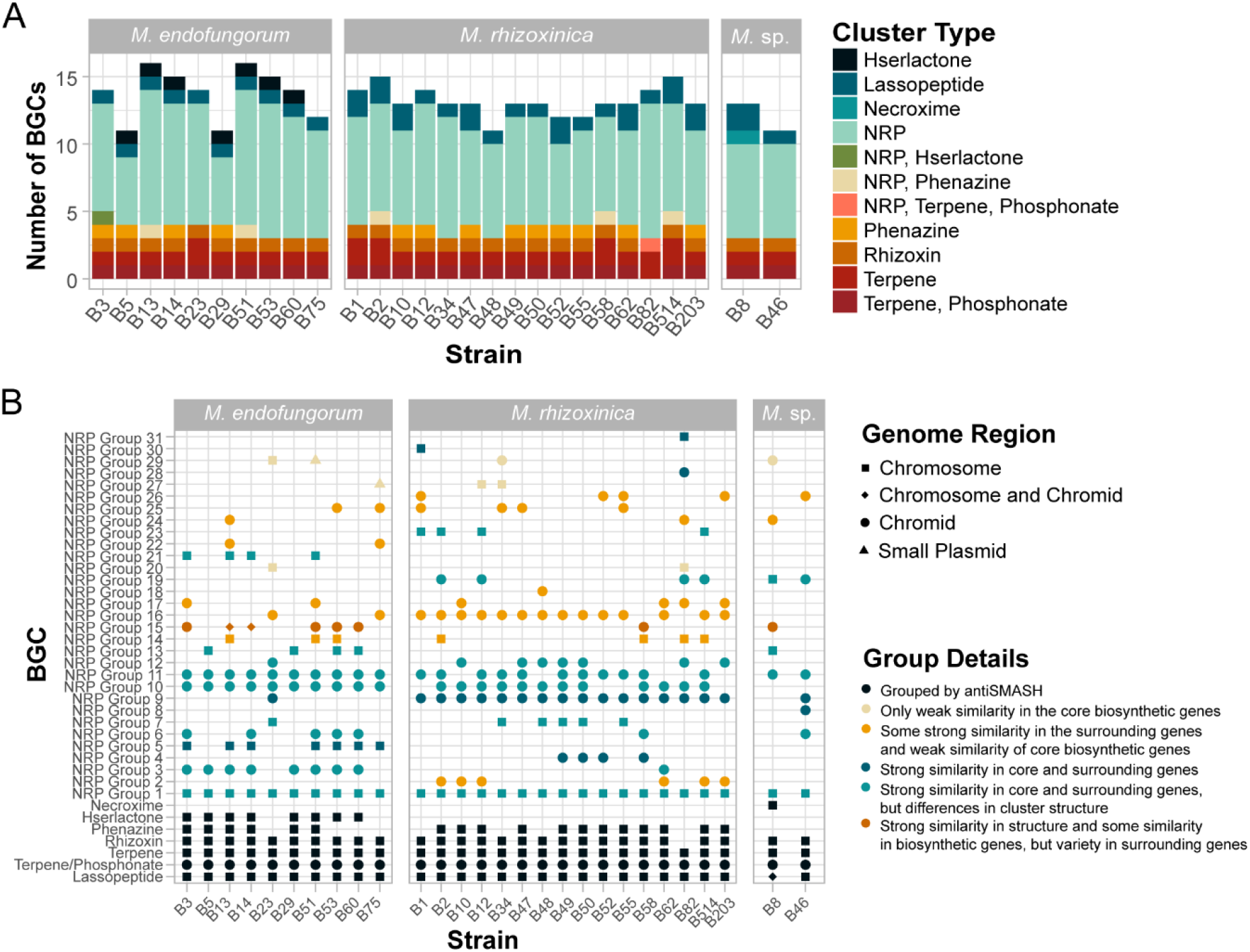
Prediction of biosynthetic gene clusters (BGCs) in *Mycetohabitans* genomes. **A**. Fractions of the types of BGC found in *Mycetohabitans* strains, grouped by species and colored by cluster type. **B**. Similarity of BGCs across *Mycetohabitans* strains and the replicons they are found in. Dots are annotated by shape and color to communicate genome region found in as well as similarity metrics. All underlying data and additional information like genomic location is in **Table S8**. NRP; nonribosomal peptide.

**Figure 6.**
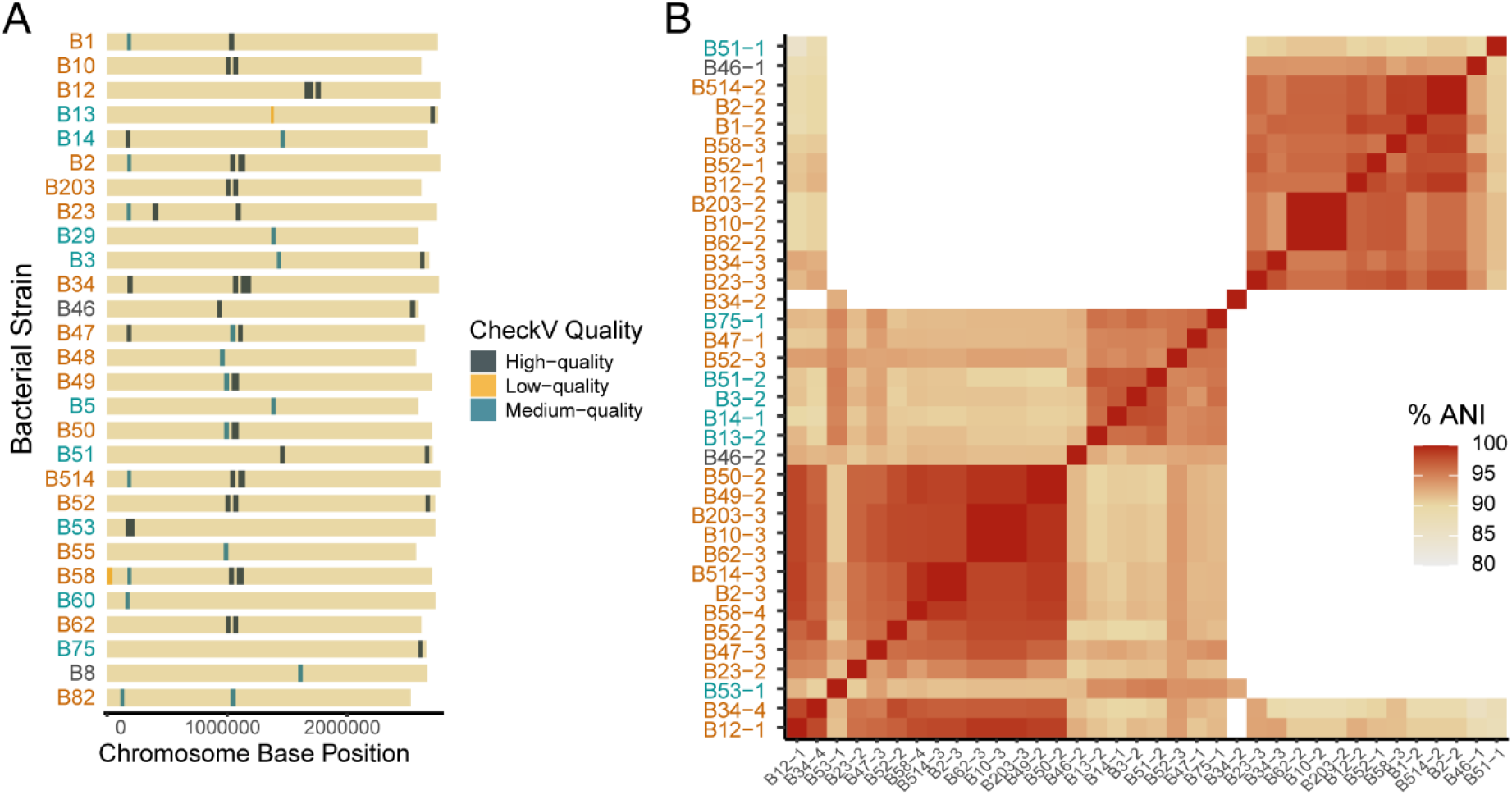
Predicted prophage regions in the *Mycetohabitans* spp. genomes. **A**. Simplified chromosome map for each strain showing the location of putative prophages indicated based on their quality score by CheckV. **B**. Average nucleotide identity (ANI) of all high-quality prophages identified across the 28 genomes, each indicated by the strain name and an identifying number. In all panels, *M. rhizoxinica* strains are indicated in orange, *M. endofungorum* in teal, and *M*. sp. in black.

We used Clinker (Gilchrist and Chooi 2021) to further group the secondary metabolite clusters beyond the initial predictions by antiSMASH, based on similarity in core biosynthetic genes and surrounding regulatory, additional biosynthetic, and transport genes. Not all NRP clusters grouped by Clinker are equally similar and clusters within a group possibly encode for different secondary metabolites. In total, there were 26 similar groups, with NRP Groups 8, 18, 28, 30, and 31 being unique/singleton BGCs. However, some BGCs are represented twice within **Figure 5B** if they were predicted to have genes for both an NRPS and another secondary metabolite product, such as phenazine. Although NRP Group 1 clusters are found across all strains, this cluster has some variety in structure with some BGCs having additional biosynthetic genes, suggesting further study is needed to assess expression and metabolite variation (**Figure S11**). NRP Group 11 clusters, found mostly in Mrh strains, have an NRPS with similarity greater than 90% to the *habA* gene (Niehs, et al. 2022). NRP Group 10 clusters are found across most strains and have core NRPS genes that are between 96-100% similar to the NRPS in the previously characterized rhizomide cluster (Wang, et al. 2018). NRP Group 9 is highly conserved and was found in all Mrh strains as well as B23 and B46. Since it does not have similarity to the NRPSs of endopyrrole A cluster nor rhizomide (**Figure S12**), this may be encoding a novel NRP of interest.

### Most *Mycetohabitans* spp. genomes have putative intact prophage

We surveyed each genome for CRISPR-Cas systems that may serve as antiphage defenses and found no *cas* operons in any strain. A subset of the strains had low-confidence, single spacer CRISPR arrays, but these had no repeat similarity between them and likely represent false positives. We thus hypothesized that the genomes may contain intact prophage. Across the set of *Mycetohabitans* spp. genomes, all were predicted to have viral sequences on their chromosomes (**Figure 5A**), but not on the chromid or plasmids, with an average per strain of 1.3 high-quality prophages.

Sixty prophage sequences were found, 36 of which were designated as high quality (predicted to be >90% complete). Seven genomes did not have any high-quality prophages: Msp B8, Mrh B82, Mrh B55, Mrh B48, Mef B5, Mef B60, and Mef B29. We ran ANI analysis to briefly assess the relationship of the high-quality prophages (**Figure 5B**). The predicted phages clustered in two distinct groups, except phage B34-2 which fell outside of both groups. Overall, Mrh tends to have a greater quantity of predicted higher quality phages, and Mef has a few strains with just viral genome fragments. Due to the novel nature of these phages, functional annotations within the prophage genes are mostly limited to standard structural components such as head proteins, tail, baseplate, etc. However, some of the genomes in the smaller of the two ANI clusters (upper right) have putative toxin/antitoxin genes and some on the middle group within the larger cluster has NtrX-like response regulators, suggesting possible functional outcomes from phage presence.

## DISCUSSION

With this study, we aimed to uncover the genome dynamics during the transition of an endosymbiotic bacterium from facultative to obligate, using the endofungal *Mycetohabitans* genus as a model. By sequencing genomes with high quality, long-read Oxford Nanopore Technology sequencing we were able to dramatically increase the genomic resources available for this genus and conduct a more structurally focused assessment. Although *Mycetohabitans* are known to have low unique gene content compared to *Burkholderia* sensu lato, we still see an open pangenome with accumulation of non-essential and/or pseudogenes, a phenomenon typically observed in genomes of recently host-restricted endosymbionts (McCutcheon and Moran 2012; Hendry, et al. 2018; Lind, et al. 2018; Bach, et al. 2022; Dewar, et al. 2024; Grujcic, et al. 2025). Added to this, the common trait of BGC enrichment resembles genome expansion of genes relevant to the role within a host, as demonstrated by insect symbiotic *Arsenophonus* spp. (Siozios, et al. 2024). In many symbionts, seemingly non-core genes contributing to host functionality are conserved despite genome loss (Carlier, et al. 2016). Overall, our pangenomic data for *Mycetohabitans* supports a hypothesis of expansion of transposable elements and BGCs, in the midst of genome reduction occurring, and likely the loss of the third replicon, though without as dramatic an expansion of pseudogenes and TEs as have been observed in other endosymbionts.

Species boundaries within endosymbiotic genera are not well delineated, generally, and require more pangenomic studies and functional studies to assess diversity and grapple with how to define species. For example, the highly studied *Wolbachia* genus is divided into various monophyletic “supergroups” with genomes that have ∼93% or higher identity within them and appear to have undergone host switching across insect species (Wang, et al. 2020; Mahmood, et al. 2023; Vancaester and Blaxter 2023). In contrast, many other endobacteria from single host species have high ANI values of >99% across genomes, such as those from the scaly-foot snail (*Chrysomallon squamiferum*) and the deep-sea mussel (*Gigantidas platifrons*) (Lan, et al. 2022; Sun, et al. 2023). Our investigation of ANI across the *Mycetohabitans* genomes revealed high intragenus identity with less clear species boundaries. As two species have been named for the *Mycetohabitans* genus, we propose conserving these species designations and setting the ANI thresholds for *Mycetohabitans* spp. at 94% to indicate species based on the observed areas of discreteness (Konstantinidis 2023). Though bacterial species are a complex concept, the assignment of this species boundary also aligns with the finding that homologous recombination efficiency decreases dramatically by 90% nucleotide identity (Power et al., 2021; Konstantinidis, 2023); though, genetic exchange within this genus has not been explicitly investigated.

Within the historical context of continued efforts to appropriately taxonomically classify members of the diverse *Burkholderia* genus, we find that our data supports *Vallotia* remaining its own genus, in contrast to the proposal to combine it with *Mycetohabitans* (Szabó, et al. 2021). Our analysis of *Burkholderia* sensu lato with all available *Mycetohabitans* and formerly *Candidatus* Vallotia genomes clearly shows that the two genera form distinct clades and have ANI separation (**Figure 1**). This conclusion is further supported by the significant genome reduction in *Vallotia* compared to *Mycetohabitans* and complete difference in niche, as all sequences assigned to *Vallotia* are from adelgids (Insecta: Hemiptera: Adelgidae), while all *Mycetohabitans* strains are fungal-associated. That these two genera are both endosymbionts that have undergone clear specialization for their unique hosts serves as an interesting natural, evolution experiment and thus could be compared for insights into signatures of endosymbiosis. Further analyses incorporating *Mycoavidus* and *Glomeribacter*, or other Burkholderia-related endosymbionts would yield more insights as well (Uehling, et al. 2017; Amses, et al. 2023).

Like many other *Burkholderia* species, *Mycetohabitans* has a multi-partite genome (diCenzo and Finan 2017; Bach, et al. 2022). Small syntenic blocks between the Mrh B1 plasmid with chromosomes and chromids of other genomes (**Figure 2A**) suggests potential recombination events between the chromosome, chromid, and plasmid, especially as the plasmid has been lost from some strains. The plasmids, being enriched in transposases (**Figure 4A**), may be functioning as a recombination hotspot, mediating gene loss (Hendry, et al. 2018). Alternatively, we hypothesize that the Type II toxin and antitoxins conserved across all *Mycetohabitans* plasmids may be the reason why the plasmids are maintained by *Mycetohabitans* spp. This could be validated by knocking out this toxin-antitoxin system and testing whether the microbe retains the plasmid.

We are considering the secondary replicon that is conserved across genomes to be a chromid, as defined in Harrison, et al. (2010). Though some guidance suggests that %GC variation between chromosomes and chromids should be less than 1% (diCenzo and Finan 2017), we observed a much higher differences between the plasmid and the chromosome (on average 4.43%) to the difference between the chromid and chromosome (on average 1.65%). Ultimately, the *Mycetohabitans* chromid has chromosome-like evolutionary dynamics including low diversifying evolution rate and a similar %GC content (**Figure 3C-D, Table S2**), as well as conservation of genes involved in core metabolism like transcription and signal transduction (**Figure S6**). Conversely, it shows plasmid-like gene content including toxin-antitoxin maintenance system, partitioning system, mobilome, and relatively high accessory gene content compared to the chromosome (**Figure 3A-B, Figure S6**).

Despite having numerous transposable elements and putative prophage, *Mycetohabitans* spp. genomes did not have any CRISPR-Cas systems, which are implicated in controlling deleterious mobile genetic elements (Westra and Levin 2020). Yet, this aligns broadly with *Burkholderia* sensu lato, as most members have orphan CRISPR arrays with no *cas* operons, or no arrays at all, though we did not survey for non-CRISPR/Cas antiphage defenses (Seo, et al. 2015; Pratama, et al. 2018). The extensive presence of toxin-antitoxin systems on the plasmids enabling their maintenance, combined with the lack of CRISPR-Cas systems may be allowing the bacteria to carry out sustained mobilization-mediated gene loss via pseudogenization (Vigil-Stenman, et al. 2015; Siozios, et al. 2024). We found that *Mycetohabitans* strains do have higher TE and pseudogene content compared to other symbiotic *Burkholderia* sensu lato. However, how many transposons are actually functional is also to be questioned (Cordaux 2009), and conclusions about pseudogenization would require further studies with data assessing transcriptional activity (Goodhead, et al. 2020). The presence of mobile genetic elements also supports horizontal gene transfer, such as type III effector acquisition and/or functionalization towards symbiosis maintenance (Dale, et al. 2001; Carter, et al. 2020; Carpenter, et al. 2024; Siozios, et al. 2024).

This repertoire of *Mycetohabitans* genomes is derived from Rm isolates sampled from across the world, and the isolates corresponding to this genomic resource are accessible through public culture collections. The majority of the samples are from United States and Southeastern Asia, which represents a strength in covering diversity from these areas but a potential gap in other regions. The orphan strains Msp B8 and B46 are the only samples from Australia and South America, respectively, which may underlie their lack of identity with the named species and supports continued sampling and sequencing to capture missing diversity. Especially given that Msp B8 is the only strain predicted to produce necroximes through a BGC, though other endosymbionts in other genera do, supporting the hypothesis of undiscovered diversity on continents in the global south (Niehs, et al. 2020; Büttner, et al. 2021). Additionally, we do not well understand the diversity represented within *Rhizopus* host fungi of *Mycetohabitans*, or the rate of horizontal transmission of symbionts. Connecting the bacterial genomic data presented here with improved fungal genomic resources and phylogenetics is a natural next step towards understanding how *Mycetohabitans* genetic diversity and speciation has been impacted by host organisms.

## METHODS

### Screening Rhizopus strains for presence of Mycetohabitans

Genomic DNA was extracted from *Rhizopus* spp. grown on Potato Dextrose Agar (PDA) overnight at 28°C using the Quick-DNA Fungal/Bacterial Miniprep Kit (Zymo). PCR using primer sets for bacterial 16S rDNA gene (27F: 5’ AGAGTTTGATCTGGCTCAG 3’; 1492R: 5’ GGTTACCTTGTTACGACTT 3’) was done to confirm bacterial presence (Heuer, et al. 1997). Fungal ITS sequence was amplified as a positive control for gDNA quality using the primers ITS86F (5’ GTGAATCATCGAATCTTTGAA 3’) and ITS4 (5’ TCCTCCGCTTATTGATATGC 3’) (Turenne, et al. 1999). Standard buffer and protocol for OneTaq 2X Master Mix (New England Biolabs) was used with 30 cycles. Products were visualized using agarose gel electrophoresis.

### DNA extraction

*Rhizopus* spp. were grown on PDA at 28°C until non-sporulating fungal mycelia was visible (2 days). Bacterial extractions were carried out by disrupting hyphal tissue suspended in 10 mM MgCl_2_ using a bead beater, micropestle, or vortex. Bacteria were separated from fungal hyphae by syringe filtration through 2-2.7 µm filters and plated on King’s medium B or lysogeny broth amended with 1% glycerol, each prepared with 1% agar and 100 µg/mL cycloheximide. After incubation at 28°C until colonies were visible, *Mycetohabitans* spp. colonies were inoculated in LB with 1% glycerol or KB broth at 28°C for 3 days. DNA extractions were done using the Wizard® Genomic DNA Purification Kit (Promega) following the protocol for gram-negative bacteria, from turbid bacterial cultures.

### Genome sequencing and assembly

The quantity of genomic DNA was assessed using a QubitFlex and sequenced by Plasmidsaurus Inc. using Oxford Nanopore Technology. Library preparation was carried out using v14 library prep chemistry with minimal fragmentation and the libraries were sequenced using Oxford Nanopore Technologies R10.4.1 PromethION flow cells. Assembly was also carried out by Plasmidsaurus using Filtlong v0.2 (github: rrwick/Filtlong) for eliminating low quality reads and downsampling when necessary, Miniasm v0.3 (Li 2016) and Flye v2.9.1(Kolmogorov, et al. 2019) for assembly generation, and Medaka v1.8.0. (github: nanoporetech/medaka) for polishing. The circular assemblies of chromosomes and plasmids were rotated to *dnaA* and *repO* respectively using dnaapler (Altschul, et al. 1990; Bouras, et al. 2024) and were submitted to NCBI under the BioProject PRJNA1167355, and annotated by PGAP (Tatusova, et al. 2016). Since the initial assembly of Mrh B62 was not circularized, the identical assembly pipeline was followed with lower stringency of downsampling to obtain a circular genome. The *Mycetohabitans* genome assemblies were analyzed for completeness and quality using BUSCO with specifying the lineage as burkholderiales_odb10, and CheckM (v1.2.2) (Parks, et al. 2015; Simão, et al. 2015).

### Phylogenetic and Average Nucleotide Identity Analysis

To compare *Mycetohabitans* genomes to *Burkholderia* sensu lato and related genera of interest, the GTDB-Tk v2.4.0 de_novo_wf command was run with default options and with taxa filters for *Mycetohabitans, Burkholderia, Paraburkholderia, Caballeronia, Trinickia, Pararobbsia, Robbsia, Mycoavidus, Glomeribacter, Pandoraea, and Chitinasiproducens* (Chaumeil, et al. 2022). *Pandoraea* was specified as the outgroup. This pipeline pulls from the current GTDB taxonomy and uses the WAG+GAMMA model within FastTree to calculate trees. In total, 383 sequences were aligned using 120 markers based on the 27 *Mycetohabitans* genomes; our B1 assembly was excluded, as the previous assembly for B1 is the reference genome. For average nucleotide identity analysis, representative genomes of each genus in *Burkholderia* sensu lato were retrieved from NCBI (**Table S4**). Together with the *Mycetohabitans* genomes, they were analyzed with FastANI (Jain, et al. 2018). For phylogenetic comparisons of chromosomes, chromids, and plasmids, replication intiation sequences were extracted from each replicon. For the chromid and chromosome, the sequences from B1 were submitted to blastn, excluding the genus *Mycetohabitans*, and the top hit was added to the analysis for an outgroup. For the plasmids, *parA* was used. Sequences were aligned with MUSCLE within MEGAX (Kumar, et al. 2018). Evolutionary history for each group was inferred with the Maximum Likelihood method and Tamura-Nei model, with default parameters and 1000 bootstraps using MEGAX. Final figures for all phylogenetic and ANI analyses were visualized and polished using FigTree v1.4.4, RStudio with key package ggtree v3.6.2 (Yu, et al. 2017), and Adobe Illustrator.

### Pangenome creation, synteny, and plasmid typing

Generation and visualization of the pangenome for 28 *Mycetohabitans* strains was done using Anvi’o v8 (Eren, et al. 2020), according to the workflow used by Delmont & Eren (Delmont and Eren 2018). *anvi-script-process-genbank* was used to split GenBank files into FASTAs, external gene calls and functions. For each *Mycetohabitans* genome, contig databases were developed by running *anvi-gen-contigs-database* – which uses Prodigal (Hyatt, et al. 2010) – on the FASTA files, with the flag *–exernal-gene-calls* specifying “NCBI_PGAP” as the gene caller. *anvi-import-functions* was used to annotate the contigs databases with the corresponding functions file. Further, *anvi-run-kegg-kofam* was run to further annotate the contig databases with KEGG metabolic pathways. The contig databases were linked using *anvi-script-gen-genomes-file* and made into a unified genomes database using *anvi-gen-genomes-storage*. Pangenome creation and visualization was done using *anvi-pan-genome*, and *anvi-display-pan* respectively, followed by *anvi-compute-genome-similarity* for FastANI (Jain, et al. 2018). The pangenome was manually binned into Core, Accessory and Unique gene clusters on the *anvi-interactive* platform based on the number of genomes each gene occurs in. Core gene clusters were found in a minimum of 28 genomes, accessory in anywhere between 2-28 genomes, and unique in a maximum of one. Identical workflows were used for creating replicon-specific comparisons after splitting the NCBI_PGAP-annotated genbanks into the three contigs using in-house scripts (github: cartercharlotte/Mycetohabitans-pangenome). Infernal version 1.1.5 was ran to investigate presence and location of various RNAs such as tRNA and rRNA in the chromid (Nawrocki and Eddy 2013). GenBank files belonging to Mrh B1, Msp B8, Mef B5, and Msp B46 genomes were used to infer synteny using syny (Julian and Pombert 2024). E-value threshold was set to 1e-10 for the synteny analyses. MOB Suite program MOB Typer was used for characterizing plasmid types (Robertson and Nash 2018).

### Heap’s law analysis

A gene presence absence matrix was generated from the computed pangenome using the command *anvi-script-gen-function-matrix-across-genomes* with specifying “NCBI_PGAP” as the *–annotation-source* and *–gene-caller*. R scripts were used to compute rarefaction curve for the data stored in matrix using the package vegan (github: vegandevs/vegan).

### Transposable Element and Pseudogene analysis

ISEScan was used to infer TE stats on the whole genome FASTA files and the *Burkholderia* sensu lato genomes listed in **Table S1** (Xie and Tang 2017). The.sum output files were then used to plot the percent genome occupied by TEs using R. Pseudogenes predicted during PGAP annotation were collated from the genbank files into a dataframe, and visualized with R (Tatusova, et al. 2016). Pearson correlation was performed on the TE and pseudogene percentages to determine whether there is a linear relationship between the two.

### dN/dS analysis

FASTA files containing DNA sequences of single-copy core genes (SCG) (Chaumeil, et al. 2022) were pulled from the chromosome and chromid genome comparisons using the command *anvi-get-sequences-for-gene-clusters* using the arguments *--report-DNA-sequences* and -*-split-output-per-gene-cluster*. These were then aligned using ClustalW (Thompson, et al. 1994). R package Ape (Paradis, et al. 2004) was used to calculate dN/dS values for each gene in each replicon, using the sequence of each gene in B1 as reference to compare with that gene sequence in the rest of the genomes. Files having non-identical gene sequences were used in these analyses, and the files that didn’t have unique sequences were excluded.

### antiSMASH and Clinker

Assembled genome FASTA files for all strains were submitted to the antiSMASH v7 bacterial version software (Blin, et al. 2023). Strictness level of “relaxed” was selected to identify both well-defined clusters containing all required parts as well as partial clusters missing one or more functional parts. All additional features of antiSMASH were run including KnownClusterBlast, MIBiG cluster comparison, Cluster Pfam analysis, TFBS analysis, ClusterBlast, ActiveSiteFinder, SubClusterBlast, RREFinder, Pfam-based GO term annotation, and TIGRFam analysis. Subsequent results were analyzed based on the cluster type predicted, organization of genes within the cluster, and the most similar known cluster. Genbank files from each cluster predicted by antiSMASH were submitted to clinker (Gilchrist and Chooi 2021) through the command line with the identity threshold between genes being the default 30%. Clinker then performed global alignments between sequences in each cluster, produced a cluster similarity matrix, created optimal ordering by hierarchical clustering, and then output visualizations using clustermap.js. The visualizations were then manually sorted through to identify clusters where the core biosynthetic genes showed similarity as well as the non-core biosynthetic genes in the cluster such as secondary biosynthetic genes, transport, and regulatory genes.

### Bacteriophage and CRISPR Prediction

The computational tool geNomad v1.11.1 (Camargo, et al. 2024) was run on the set of Mycetohabitans spp. genomes to identify prophage. Predicted phage were quality assessed with CheckV version 1.01.01 (Nayfach, et al. 2021) to estimate completeness. Average nucleotide identity analysis was run with FastANI v1.34 (Jain, et al. 2018). CRISPRCasFinder was run on each genome to detect Cas proteins and CRISPR arrays using the general clustering model (Couvin, et al. 2018).

## DATA AVAILABILITY

All genomes assembled in this work are available at NCBI and raw sequencing reads at SRA under the BioProject ID PRJNA1167355; individual nucleotide accessions are indicated in Table 1. All scripts and code used are available at github.com/cartercharlotte/Mycetohabitans-pangenome. Accession information for genomic data not generated in this study, but used for analyses, is listed in Tables S1 and S2.

## ACKNOWLEDGEMENTS

This work was supported by start-up funding from UNC Charlotte to MEC. The authors thank the UNC Charlotte EVOLVE group, Md Moinuddin Sheam (UNC Charlotte), and David Baltrus (University of Arizona) for critical discussion of data analysis, the USDA ARS NRRL for providing fungal isolates, and the UNC Charlotte University Research Computing group for computational resources.

