## Supplementary Files for "Comparative genomics reveals multipartite genomes undergoing loss in the fungal endosymbiotic genus *Mycetohabitans*"

**Comparative Genomics of Mycetohabitans - Supplementary Information**

**Table S1**. All long-read sequenced Mycetohabitans genome assemblies used in this study.

| Host Fungal Species | Host Fungal Accession | Bacterial Species | Bacterial Strain | Collection Location | Bacterial GenBank Accession |
|---|---|---|---|---|---|
| Long read PacBio Sequencing (Carpenter, et al, 2024) | | | | | |
| *Rhizopus microsporus* | CBS111563 | *M. endofungorum* | B3 HKI 0455 | Vietnam | CP062180–CP062181 |
| *Rhizopus microsporus* | ATCC52813 | *M. endofungorum* | B13 HKI 0402 | Ukraine | CP132744–CP132745 |
| *Rhizopus microsporus* | CBS112285 | *Mycetohabitans endofungorum* | B5 HKI 0456 | Mozam-bique | CP062178–CP062179 |
| *Rhizopus microsporus* | ATCC52814 | *Mycetohabitans endofungorum* | B14 HKI 0403 | Georgia | CP132741–CP132743 |
| *Rhizopus microsporus* | ATCC52812 | *Mycetohabitans rhizoxinica* | B12 | USA | CP062175–CP062177 |
| *Rhizopus microsporus* | NRRL5546 | *Mycetohabitans sp.* | B46 | Brazil | JADBGL000000000 |
| *Rhizopus microsporus* | NRRL5547 | *Mycetohabitans rhizoxinica* | B47 | Philippines | CP062173–CP062174 |
| *Rhizopus microsporus* | NRRL5549 | *Mycetohabitans rhizoxinica* | B49 | Wisconsin, USA | CP062171–CP062172 |
| *Rhizopus microsporus* | NRRL5560 | *Mycetohabitans endofungorum* | B60 | USA | CP062168–CP062170 |
| Long read ONT sequencing – This Study | | | | | |
| *Rhizopus microsporus* | ATCC62417 | *Mycetohabitans rhizoxinica* | B1 HKI 0454 | Japan | CP184277-CP184279 |
| *Rhizopus sp. strain* F-1360. | ATCC20577 | *Mycetohabitans rhizoxinica* | B2 HKI 0512 | Japan | CP176780-CP176782 |
| *Rhizopus oryzae* | NRRL62023 | *Mycetohabitans endofungorum* | B23 | Nebraska, USA | CP184774-CP184776 |
| *Rhizopus microsporus* | NRRL5551 | *Mycetohabitans endofungorum* | B51 | Philippines | CP173333-CP173335 |
| *Rhizopus liquefaciens (Rhizopus oryzae)* | NRRL 1514 | *Mycetohabitans rhizoxinica* | B514 | Unknown | CP176773-CP176775 |
| *Rhizopus microsporus* | NRRL5553 | *Mycetohabitans endofungorum* | B53 | South Africa | CP173328-CP173330 |
| *Rhizopus microsporus* van Tieghem | NRRL6255 | *Mycetohabitans rhizoxinica* | B55 | Ohio, USA | CP173325-CP173327 |
| *Rhizopus microsporus* | NRRL5558 | *Mycetohabitans rhizoxinica* | B58 | USA | CP184771-CP184773 |
| *Rhizopus americanus* | NRRL66675 | *Mycetohabitans endofungorum* | B75 | Pakistan | CP173322-CP173324 |
| *Rhizopus microsporus* | CBS 308.87 | *Mycetohabitans sp.* | B8 HKI 0404 | Victoria, Australia | CP173344-CP173345 |

| | | | | | |
|---|---|---|---|---|---|
| *Rhizopus microsporus var. oligosporus* | NRRL 2710 | *Mycetohabitans rhizoxinica* | B10 | Indonesia | CP176778-CP176779 |
| *Rhizopus oligosporus* | NRRL 6203 | *Mycetohabitans rhizoxinica* | B203 | Djakarta, Indonesia | CP176776-CP176777 |
| *Rhizopus microsporus* | NRRL 13129 | *Mycetohabitans endofungorum* | B29 | Unknown | CP173342-CP173343 |
| *Rhizopus sp.* | NRRL2934 | *Mycetohabitans rhizoxinica* | B34 | California, USA | CP173340-CP173341 |
| *Rhizopus microsporus* | NRRL5546 | *Mycetohabitans sp.* | B46 | Brazil | CP173320-CP173321 |
| *Rhizopus microsporus* | NRRL5548 | *Mycetohabitans rhizoxinica* | B48 | Illinois, USA | CP173338-CP173339 |
| *Rhizopus microsporus* | NRRL5550 | *Mycetohabitans rhizoxinica* | B50 | New York, USA | CP173336-CP173337 |
| *Rhizopus microsporus* | NRRL5552 | *Mycetohabitans rhizoxinica* | B52 | Ohio, USA | CP173331-CP173332 |
| *Rhizopus microsporus var oligosporus* | CBS 339.62 | *Mycetohabitans rhizoxinica* | B62 | Indonesia | CP178341-CP178342 |
| *Rhizopus arrhizus* | NRRL2582 | *Mycetohabitans rhizoxinica* | B82 | Ohio, USA | CP184791-CP184792 |

**Table S2.** Guanine-cytosine percent (%GC) for each replicon, and the differences between chromosome and chromid and plasmid respectively.

| Strain | Species* | %GC | | | %GC difference | |
|---|---|---|---|---|---|---|
| | | Chromosome | Chromid | Plasmid | Chromosome – Chromid | Chromosome-Plasmid |
| **B10** | Mrh | 61.484 | 59.758 | - | 1.7260 | - |
| **B12** | Mrh | 61.270 | 59.575 | 56.804 | 1.6997 | 4.470 |
| **B13** | Mef | 60.924 | 59.559 | - | 1.3644 | - |
| **B14** | Mef | 61.037 | 59.997 | 56.263 | 1.0397 | 4.774 |
| **B1** | Mrh | 61.213 | 59.705 | 57.429 | 1.5074 | 3.783 |
| **B203** | Mrh | 61.484 | 59.758 | - | 1.7260 | - |
| **B23** | Mrh | 61.254 | 59.331 | 58.424 | 1.9230 | 2.830 |
| **B29** | Mef | 61.583 | 60.017 | - | 1.5661 | - |
| **B2** | Mrh | 61.267 | 59.223 | 56.937 | 2.0437 | 4.329 |
| **B34** | Mrh | 61.308 | 59.932 | - | 1.3766 | - |
| **B3** | Mef | 60.987 | 59.467 | - | 1.5209 | - |
| **B46** | Msp | 61.445 | 59.579 | - | 1.865 | - |
| **B47** | Mrh | 61.480 | 59.769 | - | 1.7105 | - |
| **B48** | Mrh | 61.374 | 59.461 | - | 1.912 | - |
| **B49** | Mrh | 61.312 | 59.701 | - | 1.6117 | - |
| **B50** | Mrh | 61.312 | 59.703 | - | 1.6088 | - |
| **B514** | Mrh | 61.267 | 59.224 | 56.939 | 2.043 | 4.328 |

| | | | | | | |
|---|---|---|---|---|---|---|
| **B51** | Mef | 61.396 | 59.655 | 57.262 | 1.7403 | 4.133 |
| **B52** | Mrh | 61.353 | 59.498 | - | 1.8551 | - |
| **B53** | Mef | 61.191 | 60.149 | 56.615 | 1.0417 | 4.575 |
| **B55** | Mrh | 61.540 | 59.630 | 55.863 | 1.9106 | 5.676 |
| **B58** | Mrh | 61.303 | 59.790 | 55.250 | 1.5133 | 6.053 |
| **B5** | Mef | 61.583 | 60.016 | - | 1.5663 | - |
| **B60** | Mef | 61.189 | 60.282 | 56.641 | 0.9070 | 4.547 |
| **B62** | Mrh | 61.483 | 59.758 | - | 1.725 | - |
| **B75** | Mef | 61.252 | 59.589 | 57.618 | 1.6630 | 3.63 |
| **B82** | Mrh | 61.645 | 58.948 | - | 2.6970 | - |
| **B8** | Msp | 61.396 | 59.932 | - | 1.4633 | - |

**Mycetohabitans endofungorum*, Mef; *Mycetohabitans rhizoxinica*, Mrh; *Mycetohabitans* sp., Msp

**Table S3.** MOBTyper output on plasmid sequences (separate file)

**Table S4**. Reference genomes from *Burkholderia* sensu lato genera that were used in Average Nucleotide Identity analysis for Figure 1.

| NCBI ACCESSION | SPECIES |
|---|---|
| NZ_CP012981.1 | Burkholderia cepacia ATCC 25416 strain UCB 717 |
| NZ_JFHC01000001.1 | Caballeronia glathei strain DSM 50014 contig1 |
| NZ_CP047396.1 | Candidatus Vallotia cooleyia isolate 19-005_Vallotia chromosome |
| NZ_CP047399.1 | Candidatus Vallotia lariciata isolate Ad13-081_Vallotia chromosome |
| NZ_CP080504.1 | Candidatus Vallotia sp. (ex Adelges kitamiensis) isolate Adelges kitamiensis 18-608 chromosome |
| NZ_OU343031.1 | Candidatus Vallotia tarda isolate MYVALT chromosome 1 |
| NZ_ABLD01000070.1 | Paraburkholderia graminis C4D1M |
| NZ_RBZU01000001.1 | Pararobbsia silviterrae strain DHC34 scaffold1 |
| NZ_LAQU01000001.1 | Robbsia andropogonis strain ICMP2807 contig00001 |
| NZ_PTIR01000001.1 | Trinickia symbiotica strain JPY-345 Ga0139055_101 |

**Table S5.** Syny output, shared genes across *M. rhizoxinica* B1, *M. endofungorum* B5, *Mycetohabitans* spp. B8 and B46 genomes. (separate file)
**Table S6.** Anvi'o pangenome gene cluster summaries for whole genome pangenome analyses. (separate file)

**Table S7.** Percentage of genome masked by transposable element (%TE) and percentage of total coding DNA sequences identified as putative pseudogenes for each strain.

| Species* | Strain | Plasmid presence | %TE | %Total pseudogene |
|---|---|---|---|---|
| Mef | B75 | 1 | 6.36 | 9.86046512 |
| Mef | B60 | 1 | 4.53 | 7.68983269 |
| Mef | B53 | 1 | 4.44 | 8.21012897 |
| Mrh | B52 | 0 | 4.25 | 8.57409133 |
| Mef | B51 | 1 | 4.05 | 9.5209759 |
| Mrh | B82 | 0 | 3.86 | 8.63607393 |
| Mrh | B12 | 1 | 3.39 | 8.35380835 |
| Mef | B3 | 0 | 3.33 | 8.09286899 |
| Mef | B14 | 1 | 3.12 | 9.95085995 |
| Mrh | B1 | 1 | 3.08 | 8.91544118 |
| Mrh | B2 | 0 | 3.01 | 8.74365861 |
| Mrh | B514 | 1 | 3.01 | 8.77873992 |
| Mrh | B10 | 0 | 2.79 | 7.90866976 |
| Mrh | B203 | 0 | 2.79 | 7.90866976 |
| Mrh | B62 | 0 | 2.79 | 7.87557909 |
| Mrh | B34 | 0 | 2.65 | 8.58876118 |
| Mef | B13 | 0 | 2.64 | 9.96394625 |
| Mrh | B58 | 1 | 2.59 | 8.82533825 |
| Mrh | B23 | 1 | 2.43 | 8.8762984 |
| Msp | B8 | 0 | 2.31 | 8.84562842 |
| Mrh | B49 | 0 | 2.12 | 6.84976837 |
| Mrh | B50 | 0 | 2.11 | 7.86407767 |
| Mrh | B47 | 1 | 2.1 | 6.37516689 |
| Mef | B29 | 0 | 1.91 | 8.04480652 |
| Mef | B5 | 0 | 1.91 | 8.1043956 |
| Msp | B46 | 0 | 1.78 | 7.58550627 |
| Mrh | B55 | 1 | 1.73 | 8.22097994 |
| Mrh | B48 | 0 | 1.58 | 7.82219159 |
| B_cepacia_ATCC_25416 | | - | 0.59 | 1.51669641 |
| C_glathei_DSM_50014 | | - | 0.79 | 3.1949279 |
| Ca_V_lariciata_Ad13_081 | | - | 0.18 | 5.87449933 |
| Ca_V_sp_18_608 | | - | 0 | 4.63078849 |
| P_graminis_C4D1M_ctg63 | | - | 0.76 | 3.62275449 |
| P_silviterrae_DHC34 | | - | 0.24 | 1.44442487 |
| T_symbiotica_JPY_345 | | - | 0.33 | - |
| R_andropogonis | | - | 1.43 | - |

*[*] *Mycetohabitans endofungorum*, Mef; *Mycetohabitans rhizoxinica*, Mrh; *Mycetohabitans* sp., Msp. “-“ indicates that the field was not examined.

**Table S8.** Biosynthetic gene cluster summary. (separate file)

**Supplementary Figures**

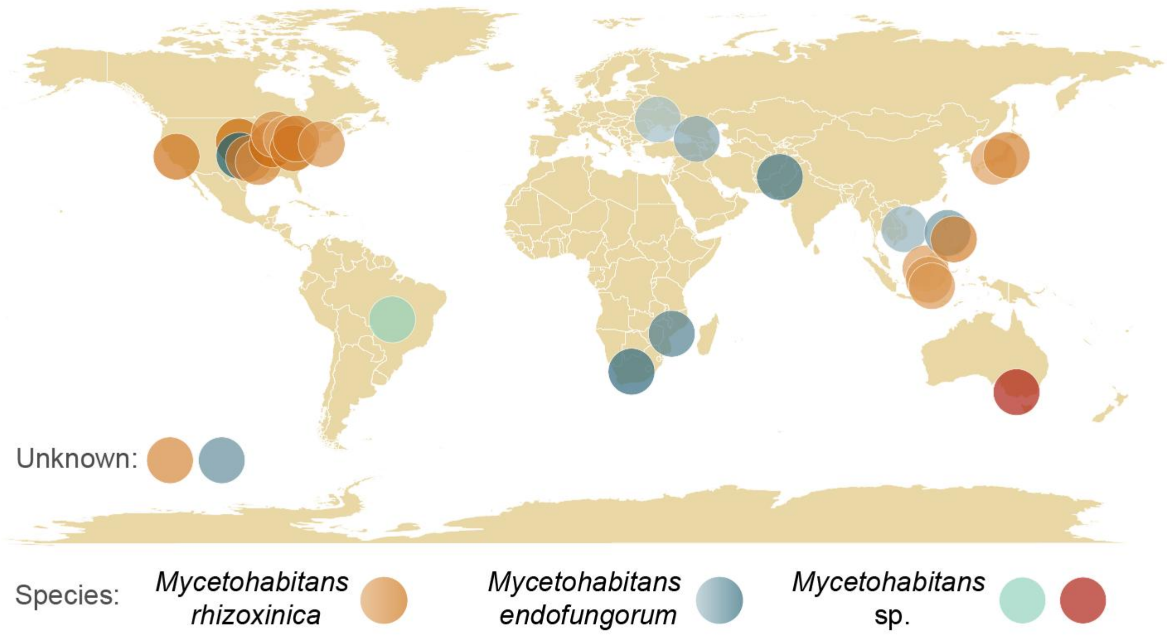


**Figure S1.** Global map showing isolation country for each fungal host of the *Mycetohabitans* strains used in this study. The species of the bacterial symbionts are indicated by color, with variation in orange and blue corresponding to strain relationship in Figure 1A, i.e. strains with similarly dark orange or blue are most closely related.

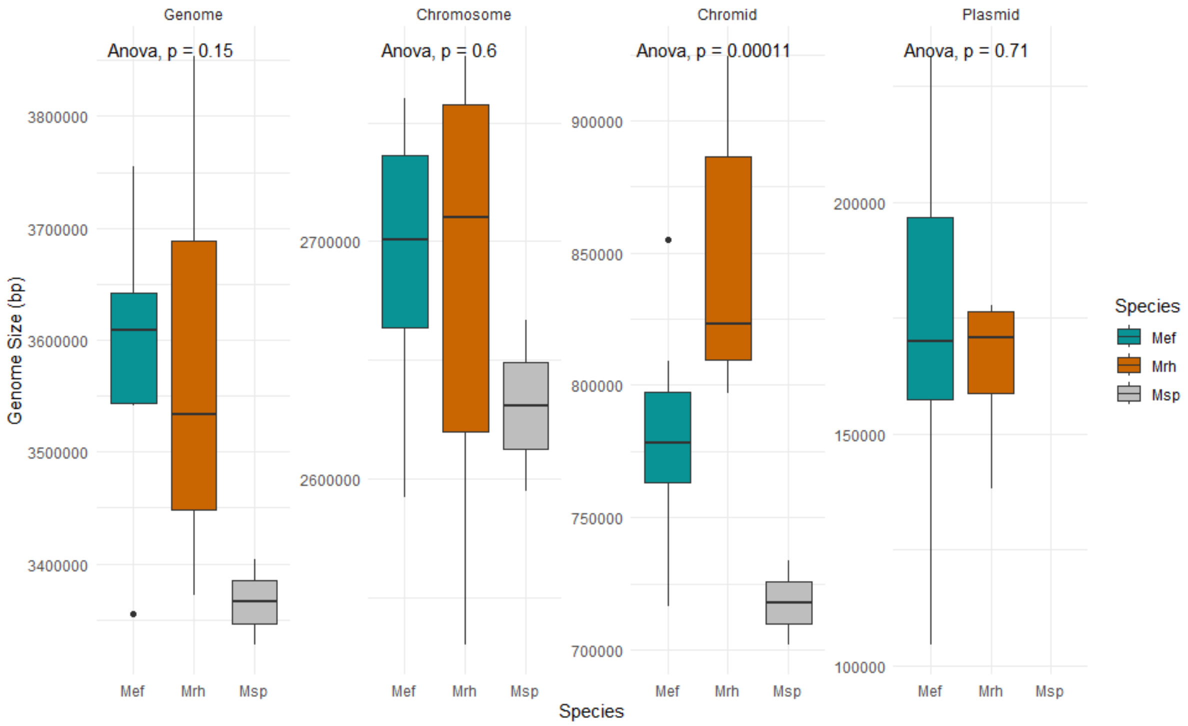


**Figure S2.** Distribution of genome and replicon sizes across all *Mycetohabitans* strains. Msp, *Mycetohabitans* spp.; Mrh, *M. rhizoxinica*; Mef, *M. endofungorum*. Means comparison was done by anova and significance value is indicated; only chromid size was significantly different by species.

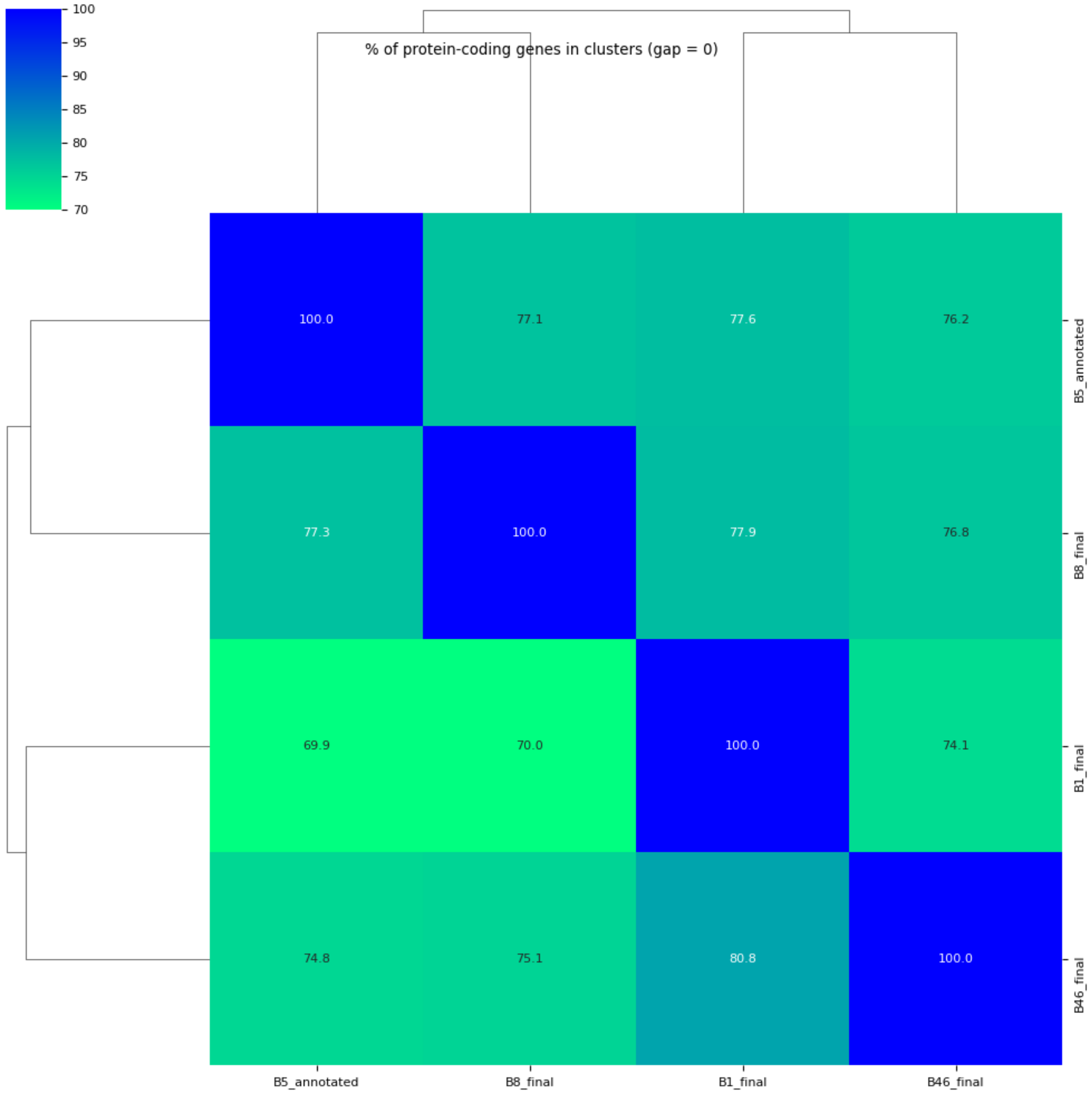


**Figure S3.** Heatmap showing hierarchical percent collinearity in genome based on protein clusters (evalue = 1e-10).

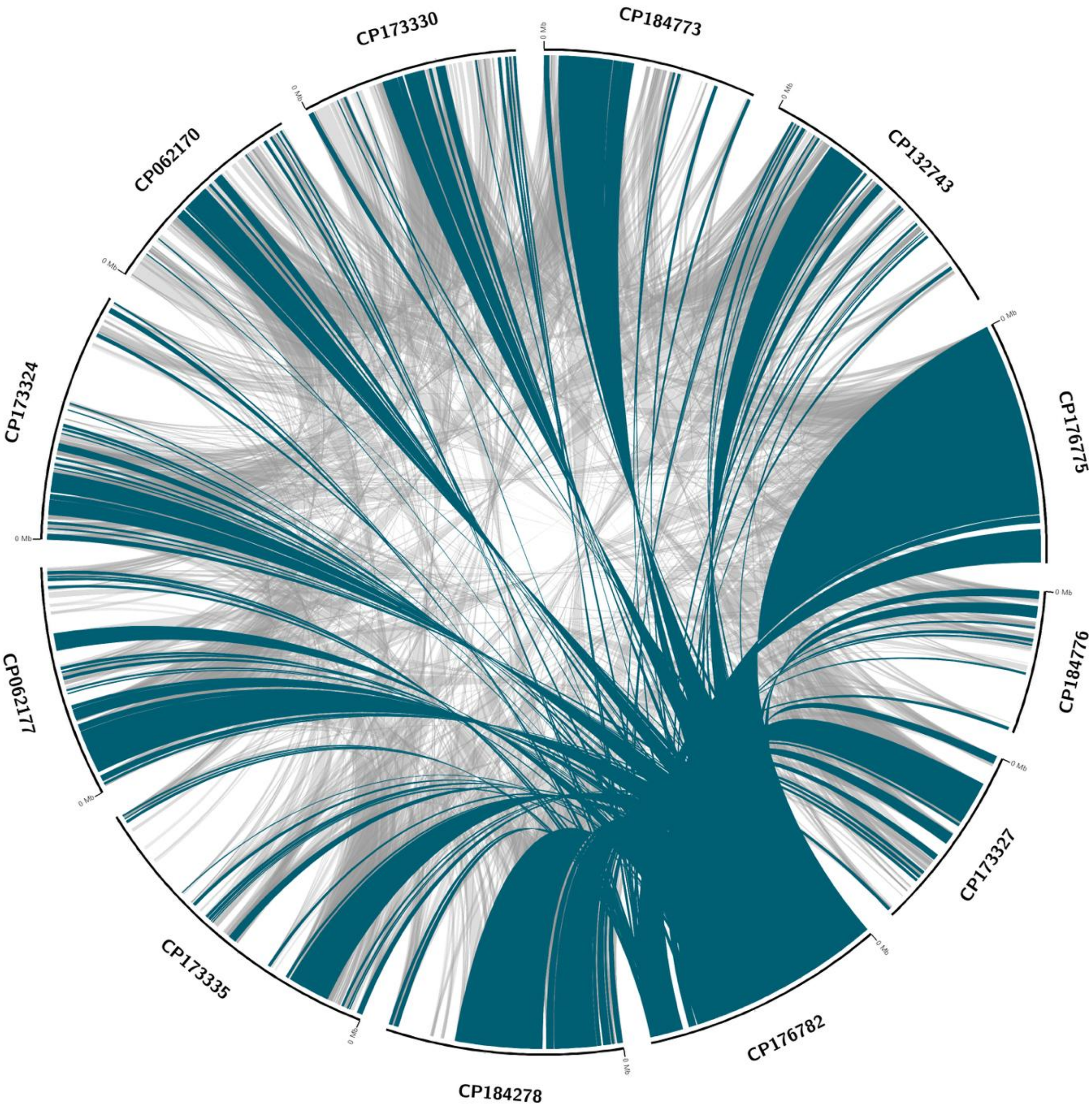


**Figure S4.** Syntenic relationship overview across *Mycetohabitans* plasmids. IDs beginning with CP correspond to NCBI sequence accessions. Teal lines indicate comparison to strain B2 as a reference.

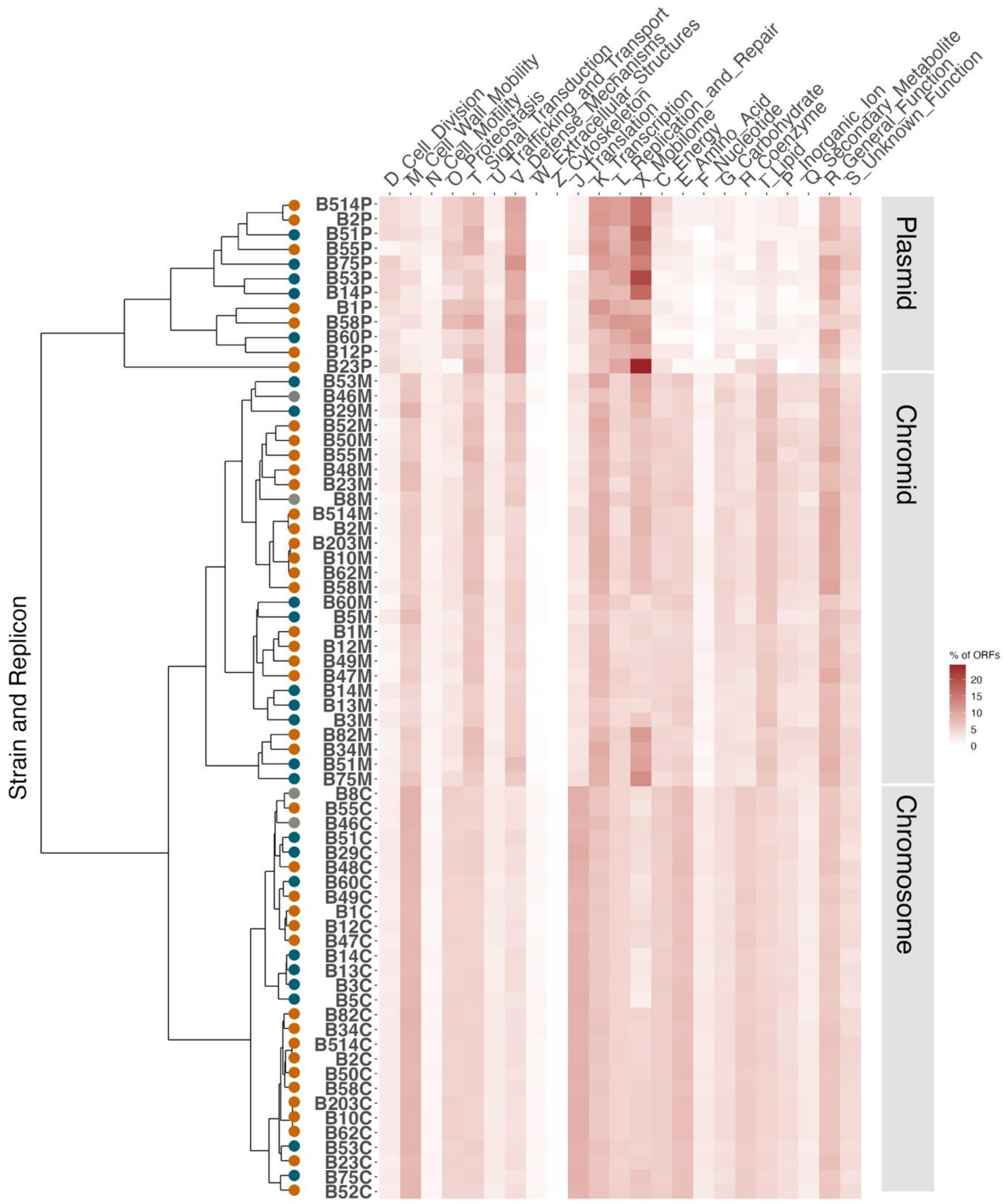


**Figure S5.** Distribution of COG gene groups distribution across replicons present in *Mycetohabitans* strains. Orange dots on the dendrogram indicate *M. rhizoxinica,* blue dots indicate *M. endofungorum*, and grey dots indicate *Mycetohabitans* sp. Chromosome, C; Chromid, M; Plasmid, P.

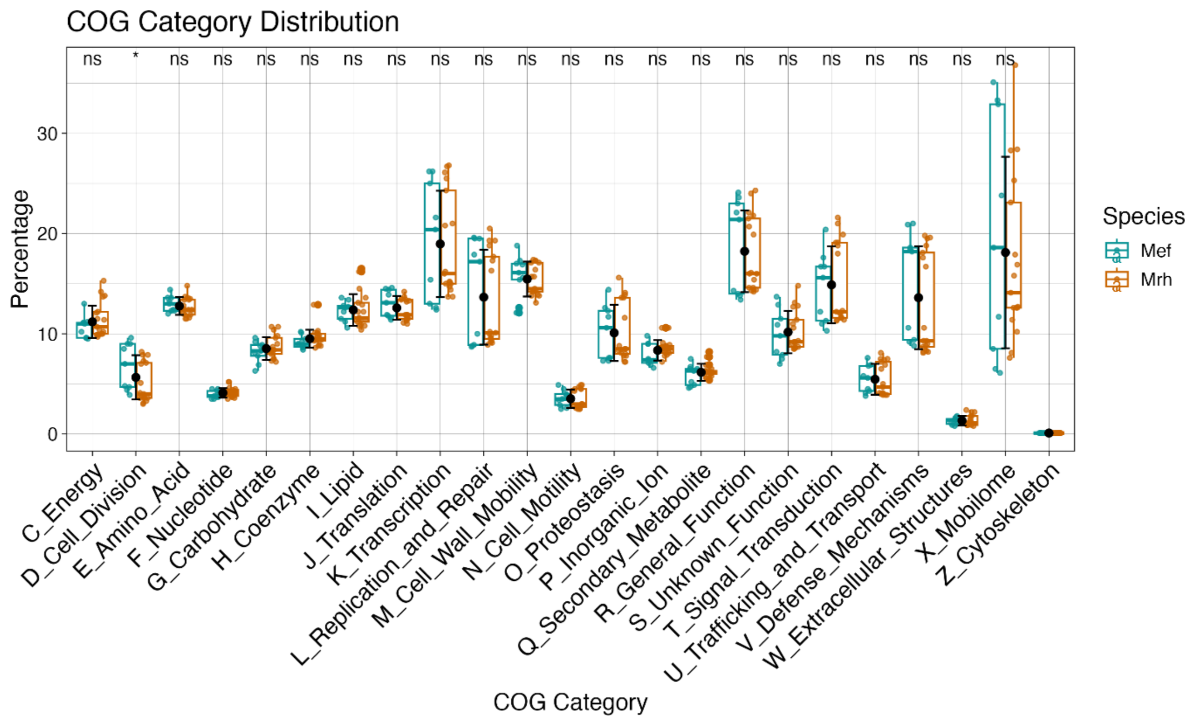


**Figure S6.** Species wise distribution of genes into functional categories. The red dot indicates the mean of all points, and the red error bar indicates standard deviation. Significance values are indicated for a Wilcoxon test between each category for the two species (p<0.05). Mrh, *M. rhizoxinica*; Mef, *M. endofungorum*.

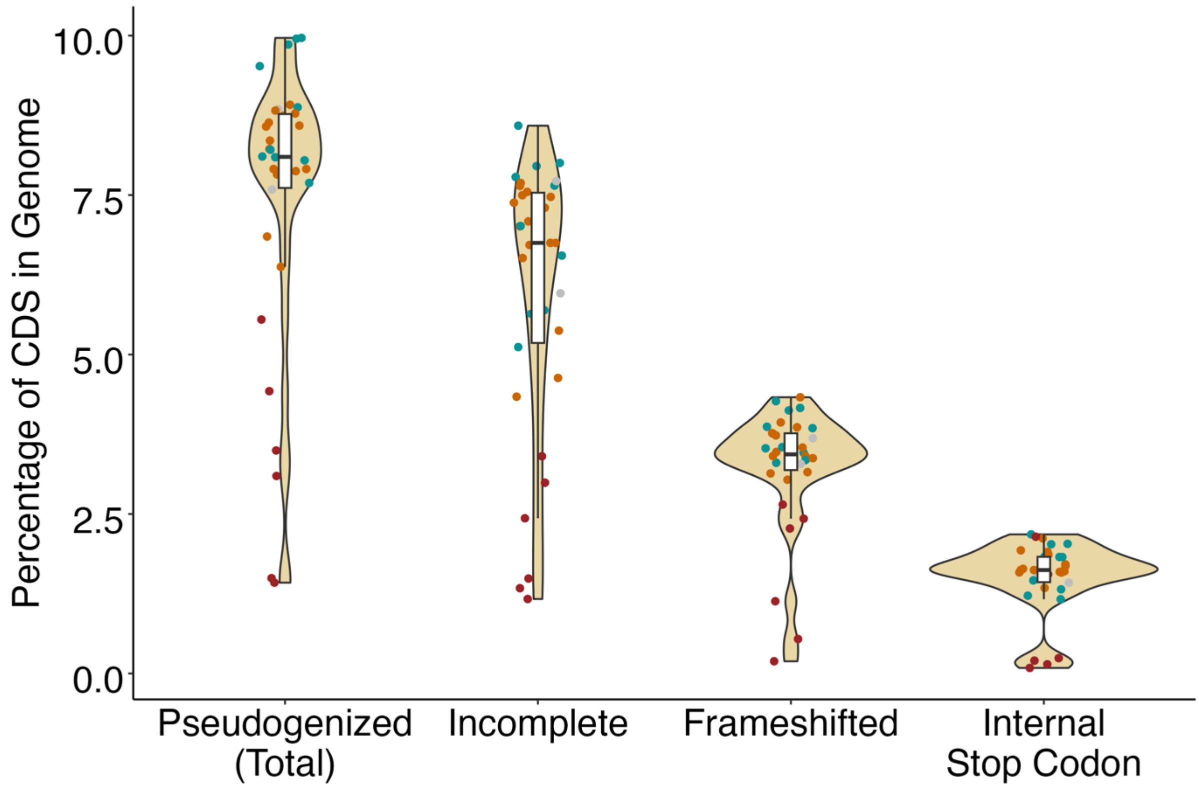

**Figure S7.** Percentage of the coding DNA sequences (CDS) putatively pseudogenized. Orange dots indicate *M. rhizoxinica,* blue dots indicate *M. endofungorum*, grey dots indicate *Mycetohabitans* spp., and deep red dots indicate representative *Burkholderia* sensu lato spp.

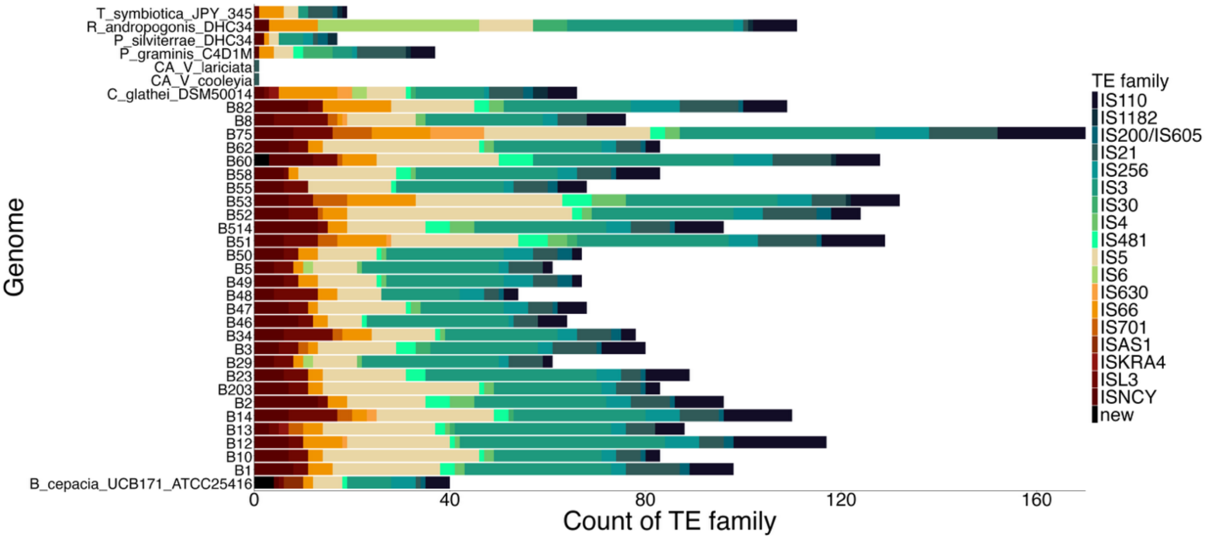


**Figure S8.** Stacked plot indicating percentage of each *Mycetohabitans* genome that is occupied by transposable elements (TEs).

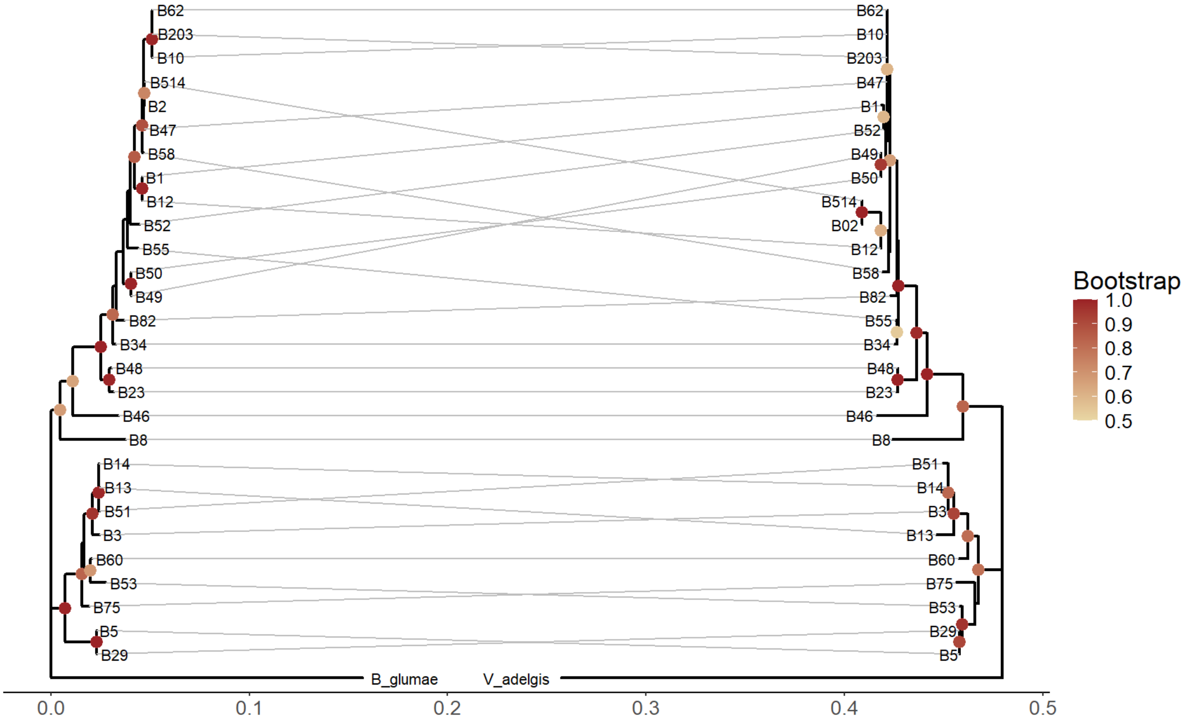


**Figure S9.** Replication origin -based phylogeny comparison between *Mycetohabitans* chromosomes (L) and chromids (R).

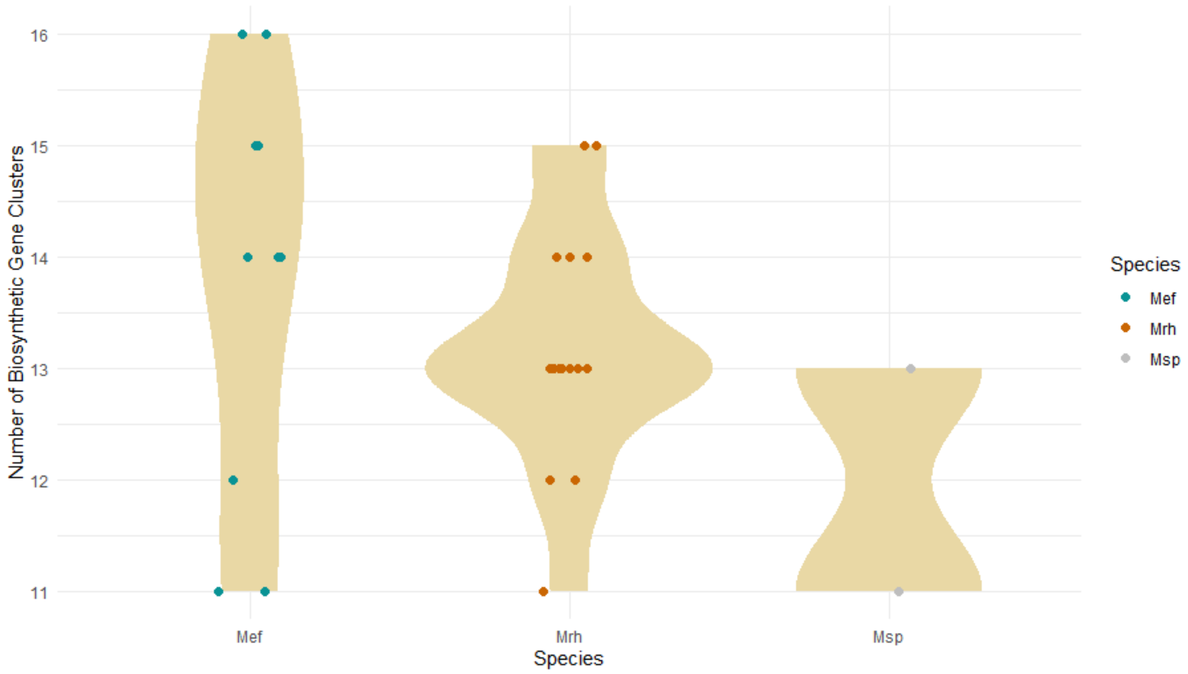


**Figure S10.** Biosynthetic gene clusters found in all *Mycetohabitans* strains. Msp, *Mycetohabitans* spp.; Mrh, *M. rhizoxinica*; Mef, *M. endofungorum*.

### antiSMASH B23

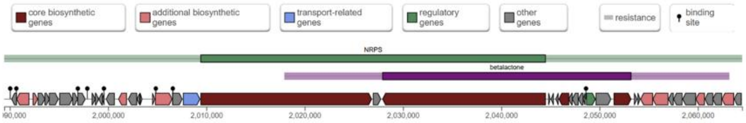


### Clinker NRP Group 1

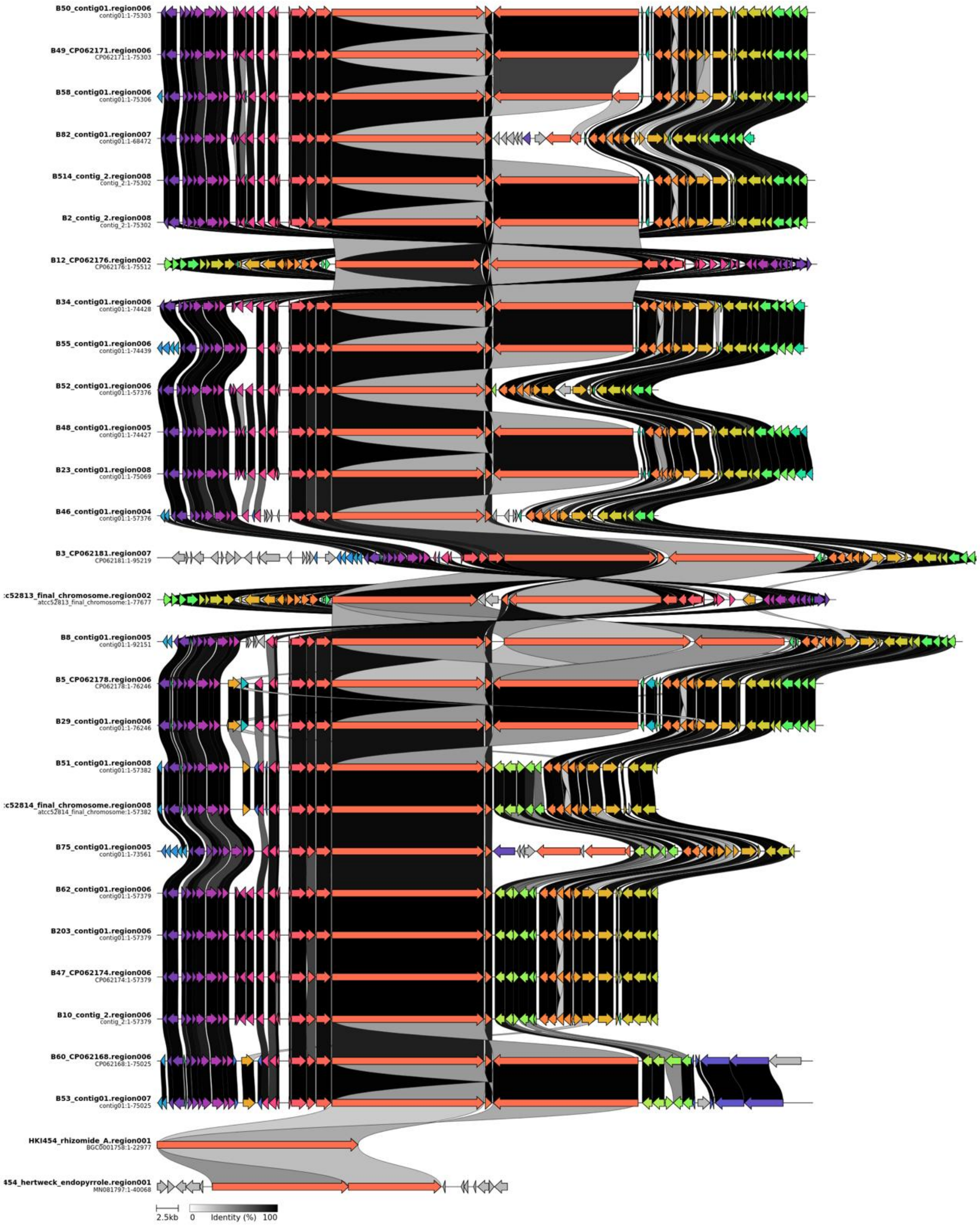


**Figure S11.** NRP Group 1 gene clusters across *Mycetohabitans* strains showing syntenic conservation and percent identity between each biosynthetic gene in the cluster between strains. Darker connectors indicate higher identity.

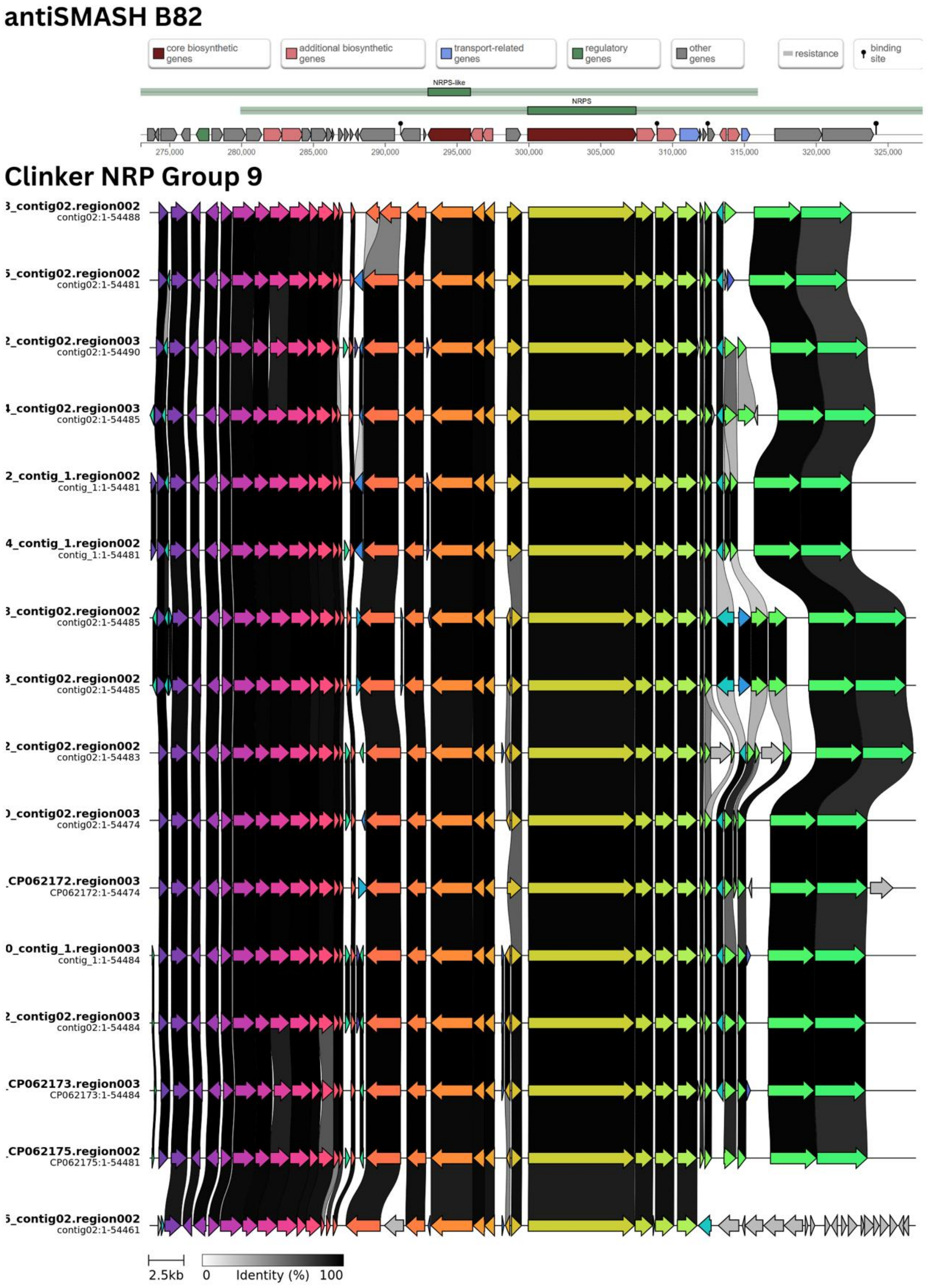


**Figure S12.** NRP Group 9 gene clusters across *Mycetohabitans* strains showing syntenic conservation and percent identity between each biosynthetic gene in the cluster between strains. Darker connectors indicate higher identity.
